# Cumulative TF binding and H3K27 Acetylation drive enhancer activation frequency

**DOI:** 10.1101/2025.03.26.645413

**Authors:** Valentina Baderna, Guido Barzaghi, Rozemarijn Kleinendorst, Kasit Chatsirisupachai, Laura Moniot-Perron, Meike Schopp, Tino Hochepied, Claude Libert, Duncan T Odom, Judith B Zaugg, Arnaud Krebs

**Affiliations:** Genome Biology Unit, EMBL, Heidelberg, Meyerhofstraße 1, 69117 Heidelberg, Germany; Faculty of Biosciences, Collaboration for Joint PhD Degree between EMBL and Heidelberg University, Heidelberg, Germany; German Cancer Research Center (DKFZ), Division of Regulatory Genomics and Cancer Evolution, 69120 Heidelberg, Germany; VIB Center for Inflammation Research, Ghent, Belgium; Department of Biomedical Molecular Biology, Ghent University, Ghent, Belgium; Structural and Computational Biology Unit, EMBL, Heidelberg, Meyerhofstraße 1, 69117 Heidelberg, Germany; University of Basel, Department of Biomedicine, Basel, Switzerland

**Keywords:** transcription factor, chromatin, nucleosome, enhancer, transcription, single molecule genomics

## Abstract

In eukaryotes, transcription factors (TFs) must continuously compete with nucleosomes to access their binding sites, leading to cell-to-cell variability in chromatin accessibility at regulatory regions. Although critical to understand enhancer function in transcription, the mechanisms that define how frequently an enhancer is active in a cell population remain unclear. Here, we used single molecule footprinting to quantify the frequency at which chromatin is accessible at enhancers in response to TF perturbations, and changes of their chromatin environment. We find that individually most TFs open chromatin in a small fraction of the cells, and that cumulative TF binding controls enhancers activation frequency. Moreover, testing the functionality of hundreds of enhancers when inserted at an ectopic genomic location identifies H3K27Ac as a chromatin mark required for their full activation. Our data support a model where enhancer activation frequency depends on the cumulative function of multiple TFs, enhanced by activating chromatin marks.

## Introduction

Gene expression is regulated by the binding of transcription factors (TFs) at *cis*-regulatory elements (CREs), such as enhancers and promoters. Activation of a CRE is a multi-step process that is influenced by both *cis*-encoded mechanisms such as TF binding and *trans*-acting mechanisms, including post-translational modification of histone tails. TFs need to open chromatin before they can stimulate transcription through the recruitment of cofactors and the general transcription machinery (1). Each CRE is bound by 5-6 TFs on average (2) and the activation of most CREs requires the collective function of multiple TFs (3–5).

Several mechanistic models have been proposed to explain how TFs combine their function to initiate and maintain chromatin accessibility. One model postulates that a specific class of TFs is specialized in opening chromatin (named ‘pioneer’), while others depend on their prior binding to exert other functions such as transcription stimulation (4, 6). In support of this concept, sequential binding and functional dependency between TFs have been observed in the activation of lineage-specifying CREs in multiple developmental contexts (4, 6, 7). Additionally, only a fraction of the TFs are able to bind their motifs when DNA is wrapped around nucleosomes, while the majority of TFs are only able to engage with free DNA (8–11). An alternative, complementary model proposes that most TFs exert only low levels of chromatin opening activity when binding individually, and that the cumulative function of multiple TFs is required to open chromatin at sufficient levels to activate transcription (4).

Chromatin accessibility is only stable for a few minutes and requires continuous maintenance through recruitment of ATP-dependent remodelers (12–14). This constant competition between TFs and nucleosomes leads to heterogeneity in CRE accessibility, where CREs are only open in a fraction of the cells at any given time (15, 16). This cell-to-cell variability in enhancer accessibility will, for instance, condition how often an enhancer can stimulate the transcription of its target gene in the cell population. Yet, the frequency at which enhancers or promoters are active in cells, as well as the molecular mechanisms involved in the regulation of their chromatin accessibility levels, are largely unknown. What is the contribution of individual TFs to enhancer accessibility frequency? How do multiple TFs combine their function at enhancers?

Each TF binding event occurs in a specific context defined by partner TFs and specific combinations of histone post-translational modifications (PTMs). The presence of PTMs is associated with activation or repression of transcriptional activity. For example, H3K27Ac is frequently used as a marker of active CREs, and presence of H3K4me1 and H3K4me3 differentiates between enhancers and promoters, respectively (17). Despite the tight association between these marks and CRE activity, the causality is not fully understood. For instance, H3K27Ac is dispensable for enhancer activity in mouse embryonic stem cells (mESCs) as the transcriptome is mostly unchanged in its absence (18). Yet, its presence has been shown to reduce the concentration of TFs required to reach transcription activation during cellular differentiation (19). The diversity of chromatin contexts encountered by TFs further complicates the quantification of their contribution to enhancer activation. To isolate the effect of TF binding on chromatin opening from the influence of chromatin context requires studying TF function under controlled conditions, at a locus where the influence of chromatin marks is minimal.

Measuring the frequency of chromatin accessibility at enhancers requires simultaneously quantifying the fraction of nucleosome-free and nucleosome-occupied DNA within a cell population. Chromatin accessibility assays such as ATAC-seq and DNase-seq are powerful tools to identify open and active regulatory elements across the genome (20). However, these assays enrich nucleosome-free DNA, losing the information about the fraction of DNA molecules that may be occupied by nucleosomes in the cell population. When performed at single-cell resolution, these assays reveal the cell-to-cell heterogeneity of CRE accessibility. Yet, only a small fraction of the genome is randomly captured in each individual cell (20, 21), making it challenging to establish the frequency at which an enhancer is open across the population. Single Molecule Footprinting (SMF) and other methylation-based footprinting assays address this issue as they sequence all DNA molecules of a sample, regardless of their chromatin accessibility state (22). The data reveal the frequency at which a CRE is accessible, occupied by nucleosomes (16, 23–25) or TFs (15, 26, 27). When sequenced at high depth, the resulting SMF data can be interpreted at the level of individual DNA molecules and used to robustly determine the frequency of occupancy states at CREs across a cell population (15, 22, 28).

Here, we leveraged SMF to measure the frequency at which chromatin is accessible at enhancers and promoters in mESCs and quantified the respective contribution of TF binding from the chromatin environment to their accessibility. We developed and used *FootprintCharter* (*unpublished*), an unsupervised method to quantify the frequency of chromatin accessibility on individual DNA molecules genome-wide. We found that the frequency of chromatin accessibility is highly variable at active CREs, with promoters often accessible across the majority of the cells in a population, while most enhancers are only open in a minority of cells at any given time. We combined genome-scale genetic perturbation of individual TF motifs, and rapid TF degradation to test the contribution of individual TFs to the frequency of enhancer activation. We found that most individual TFs contribute to a partial increment in chromatin accessibility, increasing the frequency of chromatin accessibility in less than 10% of the cells in average at any given time. Consistently, we found that the frequency of accessibility observed at most enhancers requires cumulative binding of multiple TFs at the locus. To disentangle the effects of chromatin context from TF function, we measured the capacity of hundreds of enhancers and promoters to maintain their chromatin accessibility when inserted at an ectopic locus lacking chromatin modifications. Most enhancers only partially recapitulated the frequency of chromatin accessibility measured at their native loci. We found that enhancers ability to re-establish accessibility was inversely correlated with the level of the activating chromatin mark H3K27Ac at its endogenous location. By inhibiting the histone acetyltransferase p300, we demonstrated that H3K27Ac is required to enhance the frequency of chromatin accessibility at enhancers genome-wide. Finally, we show that changes in the frequency of chromatin accessibility at enhancers directly affects their activity through reduced recruitment of RNA Pol II (Pol II).

## Results

### Chromatin is accessible in a minority of the cells at active enhancers

In contrast to bulk chromatin accessibility assays such as ATAC-seq or DNase-seq (Figure 1A), single-molecule footprinting (SMF) captures the cell-to-cell heterogeneity in the chromatin accessibility of *cis*-regulatory elements (CREs) across individual DNA molecules (Figure 1B). To quantify the frequency of chromatin accessibility at CREs, we developed an unsupervised molecular classifier and footprint detection algorithm named *FootprintCharter* (*unpublished*). The method partitions DNA molecules based on their chromatin accessibility patterns and distinguishes transcription factor (TF) and nucleosome footprints based on their width, without prior assumptions on the binding of TFs at the locus (Figure 1C, Supplementary Figure 1A).

**Fig. 1.**
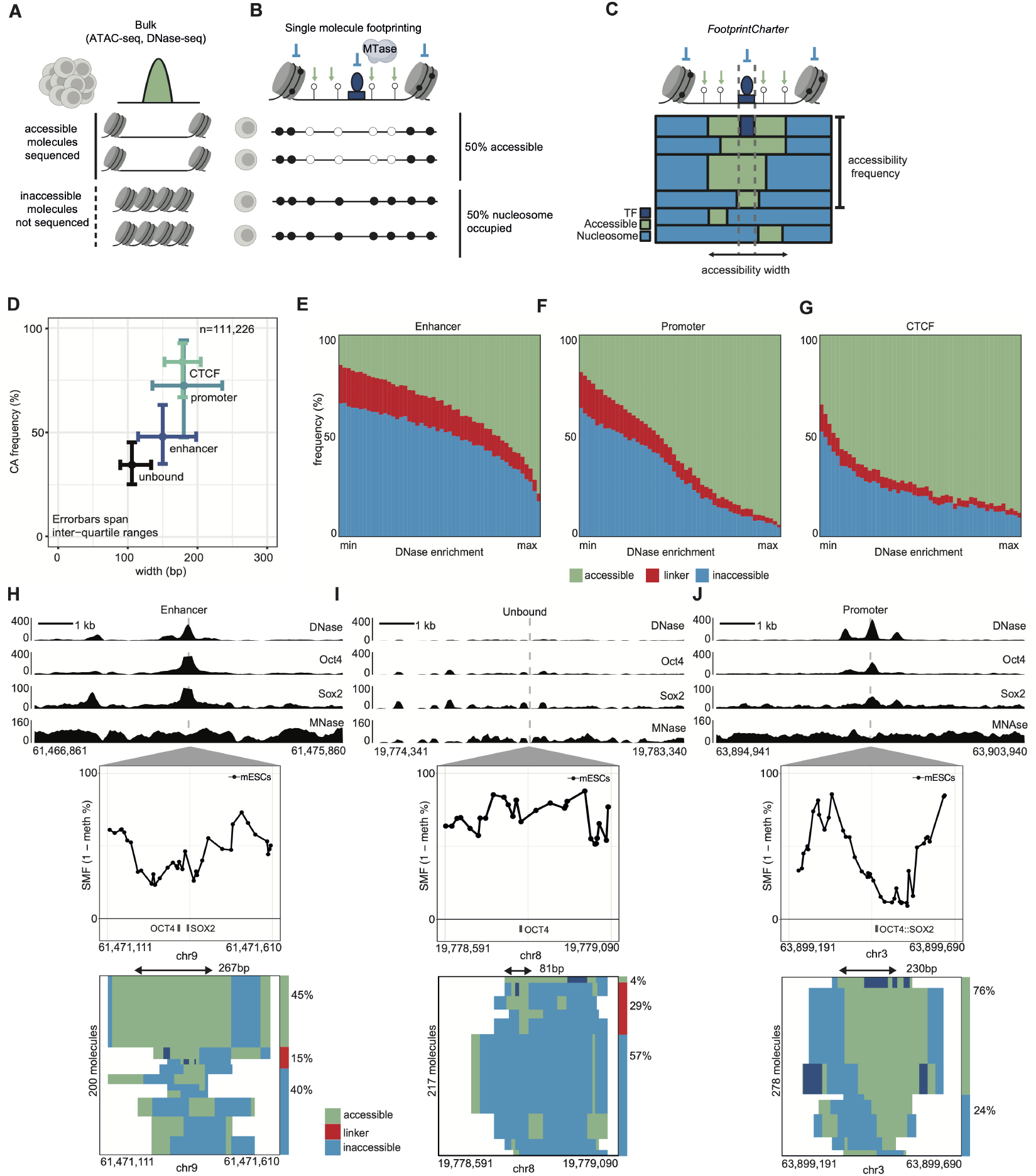
Quantification of the frequency of chromatin accessibility at cis-regulatory elements. **(A)** Mammalian CREs are characterized by different levels of accessibility. Bulk assays, such as DNase-seq and ATAC-seq, classify and rank open regulatory elements but provide only relative enrichment of the accessibility signal, limiting their ability to estimate CREs’ underlying frequency of accessibility across a cell population. **(B)** Single molecule footprinting (SMF) uses exogenous methyl-transferases to footprint protein-DNA contacts. Profiling DNA methylation by bisulfite sequencing captures and quantifies inter-cellular heterogeneity in chromatin accessibility at CREs with single-DNA-molecule resolution. **(C)** *FootprintCharter* employs unsupervised partitioning to locate and quantify footprints for TFs and nucleosomes in SMF data. Here, we use *FootprintCharter* to quantify the frequency of chromatin accessibility (CA) at TF motifs and the width of accessible stretches, averaged across molecules. **(D)** Different CREs exhibit distinct chromatin accessibility frequencies and widths. Scatter plot showing distributions of total CA frequency (y-axis) and average accessibility width across molecules (x-axis) for bound motifs at CTCF-containing loci, promoters, enhancers, and unbound motifs at intergenic regions. Dots represent median values, while error bars span interquartile ranges (IQR, 1st to 3rd quartiles). **(E-G)** Different CREs have varying frequencies associated with linker DNA, accessibility, and inaccessibility. Bar plots show distributions of molecular state frequencies (large accessible stretches in green, linker-length accessible DNA in red, inaccessible in blue) for active enhancers (**E**), promoters (**F**), and CTCF-containing loci (**G**) as a function of DNase enrichment. The accessible fraction includes molecules displaying TF footprints. CREs were binned by DNase enrichment, and the median frequency was calculated for each state and bin. **(H)** Example of an active enhancer harboring an OCT4 and a SOX2 motif. At the SOX2 motif, chromatin accessibility is detected in (60%) of the cell population, with (15%) of molecules showing accessibility widths compatible with linker DNA (<100 bp) and (45%) displaying large accessibility stretches (>100 bp). The upper panel shows genomic tracks for DNase-seq, ChIP-nexus for OCT4 and SOX2, and MNase-seq around the TF motifs. The middle panel shows the average SMF signal (1 - methylation %), along with TF motif annotation. The lower panels show stacks of single molecules sorted by decreasing accessibility widths, with chromatin states annotated (accessible in green, linker DNA in red, inaccessible in blue). *FootprintCharter* annotations indicate the average accessibility width (top arrow-headed segment) and frequencies of accessibility in active molecules (green bar) and linker DNA (red bar). **(I)** Example of an intergenic locus harboring an inactive, unbound OCT4 motif, accessible in 33%) of the cell population, primarily associated with linker DNA (29%). Same representation as in panel (**H**). **(J)** Example of an active promoter harboring a composite OCT4::SOX2 motif, accessible in 76% of the cell population exclusively in large accessibility stretches.

We used *FootprintCharter* to systematically quantify the width and frequency of chromatin accessibility around TF binding motifs (Figure 1C) in mouse embryonic stem cells (mESCs). We previously showed that TFs create footprints in SMF data, a feature resolved by *FootprintCharter*. However, footprints are not observed for all TFs and depend on the presence of cytosines in proximity to their binding sites (15, 29). To treat all TFs equally in this study, we classified molecules only based on the presence or absence of nucleosome footprints, categorizing molecules with TF footprints as accessible (see Methods). We used a previously generated bait-capture SMF dataset that profiles more than 60% of active enhancers and promoters at sufficient coverage for single-molecule quantification (15) (Supplementary Figure 1A-C). We mapped all motif instances in the genome using data from Mathelier et al. (30) and identified bound motifs using publicly available ChIP-seq and ChIP-nexus datasets as a reference for the binding of 18 TFs expressed in mESCs (Supplementary Table 1, see Methods). We first compared the distribution of the frequency of accessibility around TF motifs at various types of CREs (Figure 1D). We observed that TF motifs contained within CREs such as promoters or CTCF-bound loci showed pervasive chromatin accessibility across cells, with frequencies of accessible molecules exceeding 70% over median stretches of 180 bp and 179 bp, respectively (Figure 1D). In contrast, TF motifs within enhancers were typically accessible in only half the molecules in the cell population (48% median – blue, Figure 1D) and over shorter stretches (150 bp median). We observed that unbound TF motifs showed high levels of chromatin accessibility, with a median frequency of 35% of the molecules (Figure 1D). Yet, at these unbound motifs, we mainly observed short stretches of accessibility of about 100 bp (Figure 1D). These short stretches are consistent in length with the exposure of the TF motifs at the linker between nucleosomes (16). Thus, for subsequent analyses, we distinguished short stretches of accessibility that may correspond to linker DNA (linker-length accessible DNA) from larger accessibility stretches presumably created by the binding of TFs. This stratification by the width of accessibility confirmed the heterogeneity at active enhancers (Figure 1E) and highlights the contrast with promoters or CTCF-bound regions (Figure 1F-G). For enhancers, typically less than 50% of cells had accessible chromatin, and a substantial portion arose from linker-length accessible DNA (14% median, Figure 1E, Supplementary Figure 1D). Linker-length accessible DNA was also detectable at promoters and CTCF-bound sites, albeit at lower frequencies (Figure 1F-G, Supplementary Figure 1D). The heterogeneity in the width of chromatin accessibility at enhancers was further evident when looking at an individual enhancer bound by SOX2 and OCT4 (Figure 1H). Here, 45% of the cells showed accessible motifs associated with continuous stretches of 267 bp. The motifs on the remaining molecules were either covered by nucleosomes (40%) or accessible through linker-length stretches (15%). The same short stretches could also be observed at an example of an OCT4 motif where no binding was detected by ChIP-seq (Figure 1I), further suggesting that these correspond to the linker between nucleosomes. This is in contrast with the promoter of the *Plch1* gene, where 76% of the molecules showed accessibility stretches of *>*100 bp (Figure 1J). Overall, these data suggest that most enhancers are open for activation in less than half of the cells of a population at any given time.

### Frequency of chromatin accessibility correlates with the number of bound TFs

We next asked if binding of specific TFs, or their combination, could explain the differences in the frequency of chromatin accessibility observed between different CREs. We compared the distribution of the frequency of chromatin accessibility at the binding motifs of TFs for which we detected binding by ChIP. Only five of the tested TFs were associated with a high frequency of chromatin accessibility across their binding sites (median >70%) (Figure 2A). These include zinc-finger domain-containing TFs that are known to bind individually in the genome (CTCF, REST, and BANP), but also the histone-fold domain factor (NFY) and the general transcriptional activator NRF1. In contrast, the binding of most TFs was associated with chromatin accessibility in a minority of the molecules (22%-54%, Figure 2A). These include well-characterized regulators of pluripotency such as OCT4 and SOX2, which are critical for chromatin accessibility in this system (31, 32). Since the binding of most individual TFs was associated with partial chromatin opening, we next asked whether loci with multiple bound TF motifs would display higher proportions of molecules with accessible chromatin. We then stratified the frequency of accessibility by the number of TF binding sites that we detected to be bound by ChIP-seq at the CRE. We observed that both the frequency of accessible molecules and the width of accessibility increased with the number of TF motifs (Figure 2B, C). This effect is also observed when focusing on a specific TF, such as KLF4 (Figure 2D). TFs that are associated with a higher frequency of chromatin accessibility, such as NRF1, also showed cumulative effects but reached saturation at a lower number of binding sites, suggesting that these TFs may contribute more strongly to chromatin accessibility (Figure 2E, Supplementary Figure 2A). An example of this gradual increase in molecular frequency and width of chromatin accessibility could be observed when inspecting individual enhancers containing 1, 3, or 7 TF motifs (Figure 2F-H). Together, this suggests that individual motif instances for most TFs, including pioneer TFs, are associated with partial opening of the chromatin, and that cumulative binding of TFs may explain the frequency of chromatin accessibility observed at CREs.

**Fig. 2.**
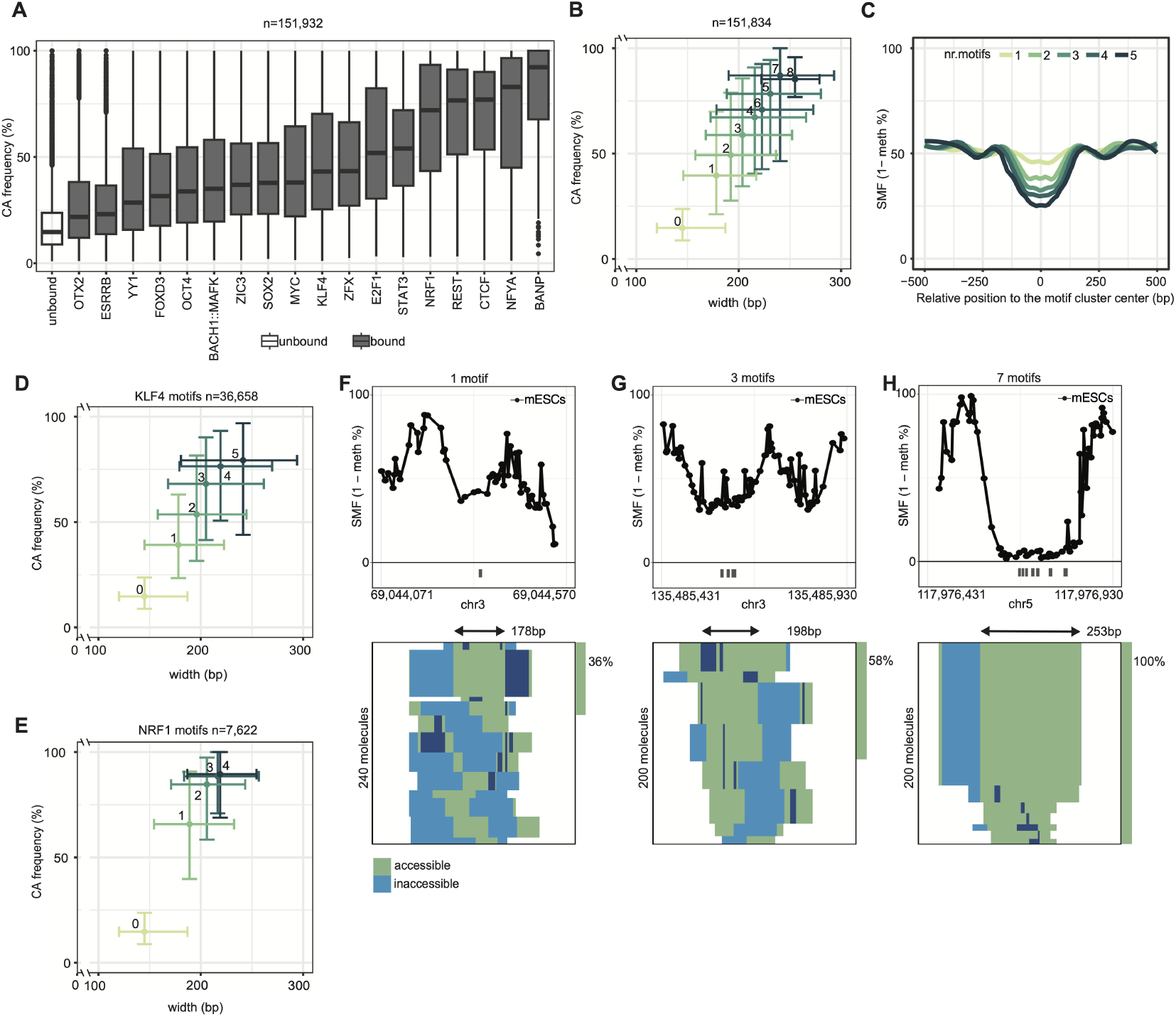
Frequency of chromatin accessibility correlates with the number of TFs bound at the CRE. **(A)** Frequencies of chromatin accessibility associated with binding of specific TFs. Boxplot representing the distribution of frequency of chromatin accessibility at motifs that are unbound (white) or bound by specific TFs (grey), as defined using ChIP-seq data. TFs with fewer than 30 quantified motifs are excluded, resulting in 151,932 total motifs reported. **(B)** Frequency and width of chromatin accessibility scale with the number of TF binding motifs at the CRE. Scatter plot showing the distributions of accessibility frequency (y-axis) and average width (x-axis) at active molecules as a function of the number of active motifs at CREs (color scale). Dots represent median values, and error bars span interquartile ranges (IQR, 1st to 3rd quartiles). TF cluster sizes with fewer than 100 data points are excluded, resulting in 151,834 total motifs reported. **(C)** Composite profile of SMF signal as a function of the number of active motifs at CREs. This representation shows the SMF signal (1 - methylation %)) for individual cytosines, averaged and smoothed across heterologous CREs. **(D-E)** Analogous representation as (**B**) but focusing on the bound motifs of (**D**) KLF4 and (**E**) NRF1. **(F-H)** Single locus examples of enhancers with (**f**) one, (**g**) three, or (**H**) seven TF-bound motifs, showing increasing frequencies of chromatin accessibility. Same representation as Figure 1H.

### The contribution of TFs to chromatin accessibility is partial and cumulative

In order to test this hypothesis, we measured the consequence of the perturbation of TF binding on the frequency of chromatin accessibility at CREs. If chromatin accessibility is the result of the cumulative binding of multiple TFs, we would expect a partial reduction in the frequency of accessible molecules upon deletion of individual motifs, and that this reduction would scale with the number of removed motifs. To quantify the contribution of individual TFs to the frequency of chromatin accessibility, we leveraged the natural genetic variation between different mouse species. Specifically, we studied the impact of single nucleotide variations (SNVs) that reduce TF motif affinity on the frequency of chromatin accessibility at the locus. We performed SMF in two F1 lines derived from crosses between *Mus musculus domesticus* and *Mus musculus castaneus* (Bl6xCast), as well as *Mus spretus* (Bl6xSpret) (Figure 3A). These cell lines have a median of 3 and 5 SNVs per CRE for Bl6xCast and Bl6xSpret, respectively (Supplementary Figure 3A). Chromatin accessibility is measured in the same nuclear environment for both alleles of each genotype, thus controlling for possible trans-effects, such as changes in TF concentrations that may occur when comparing cell lines from different species. We sequenced samples at sufficient depth to reproducibly quantify the frequencies of chromatin accessibility on each allele (Supplementary Figure 3C-I). To test the effects of deleting single motif instances, we focused on SNVs that led to strong reductions in predicted motif affinity on the alternative allele (>50%, see Methods). To increase statistical power, we jointly analyzed the SNVs occurring between the Bl6 and the Cast or Spret genotypes. We bench-marked our approach by quantifying changes in DNA occupancy across alleles using CTCF, which mostly binds in isolation from other TFs and for which single motif instances are sufficient to drive chromatin opening (9). We indeed observed strong allelic changes in frequency of accessibility for loci that harbor an SNV that strongly affects the affinity of the CTCF motif (Supplementary Figure 3J), while loci with unaltered motifs show no changes in chromatin accessibility between alleles (Supplementary Figure 3K-L). This confirms the sensitivity and specificity of the single-molecule quantification of chromatin accessibility across alleles.

**Fig. 3.**
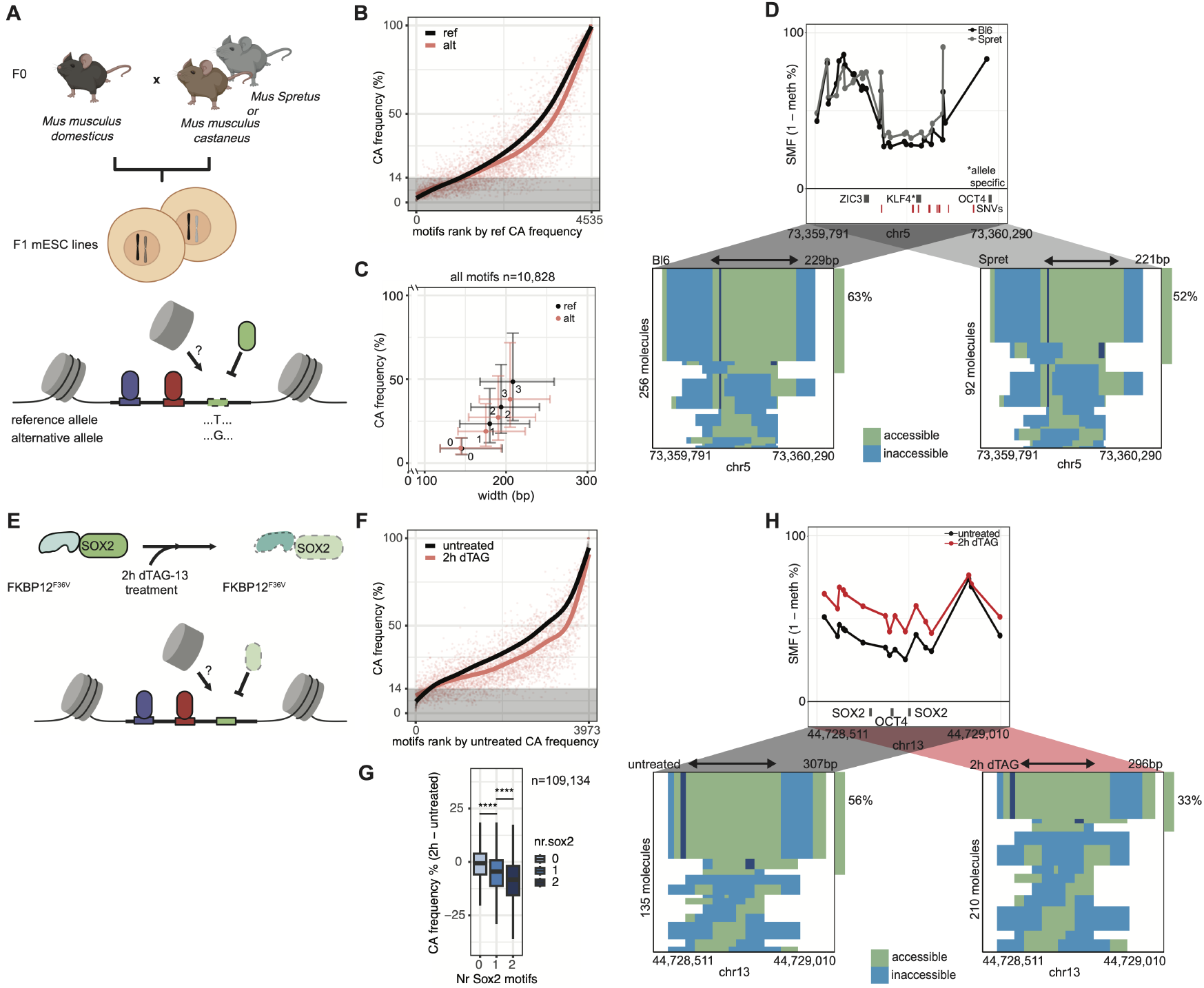
Frequency of chromatin accessibility requires cumulative function of multiple TFs. **(A)** Schematic representation of the experimental strategy used to achieve systematic genetic perturbation of TF binding. We leveraged genetic variation between evolutionarily distant mouse species and generated hybrid F1 mESCs. The two alleles in the F1 mESCs share and are exposed to the same nuclear environment, eliminating possible trans-effects. We performed SMF in F1 mESCs derived from two different crossings and measured the allele-specific frequency of chromatin accessibility for TF motifs with binding affinity disrupted by SNVs on the alternative allele (the T on the reference allele is a G on the alternative allele). The disrupted affinity could abolish TF binding and promote nucleosome occupancy. **(B)** Most TF disruptions lead to small changes in the frequency of chromatin accessibility at CREs. Scatter plot of chromatin accessibility (CA) frequencies in the reference allele (black) and the alternative allele (red) at loci where an SNV disrupts a TF motif. The shaded area indicates the background accessibility observed in closed chromatin. **(C)** Loss of individual TFs shifts the frequency of chromatin accessibility to predictable levels. Scatter plot showing allele-specific frequency of accessibility (y-axis) and average width of accessibility (x-axis) at active molecules as a function of the number of bound motifs at CREs (reference and alternative alleles are shown in black and red, respectively). Dots represent median values, and error bars span interquartile ranges (IQR, 1st to 3rd quartiles). **(D)** Example locus of an enhancer bound by three TFs, including a KLF4 with a strong difference in affinity between the Bl6 and Spret alleles. This genetic variation is associated with an 11% reduction in chromatin accessibility frequency in the cell population. The upper panel shows the average SMF signal (1 - methylation %) for the BL6xSpret F1 line (black dots and line for Bl6, grey dots and line for Spret), as well as the TF motifs and SNV annotations. The lower panels show *FootprintCharter* stacks for the Bl6 (left) and Spret (right) alleles. **(E)** Schematic representation of the experimental strategy used to test the impact of deleting individual TFs. To this end, we used rapid degradation of SOX2 via the dTAG system. The tag FKBP12^*F* 36*V*^ is fused to SOX2. By adding the degrader molecule dTAG, the chimeric protein is ubiquitinated and degraded by the proteasome. **(F)** Loss of SOX2 leads to small changes in the frequency of chromatin accessibility at CREs. Scatter plot of chromatin accessibility frequencies in untreated cells (black) and cells treated for 2h with dTAG against SOX2 (red) at SOX2-bound loci. The shaded area indicates the background accessibility observed in closed chromatin. **(G)** A higher number of SOX2 motifs at the CRE increases the frequency of chromatin accessibility loss upon SOX2 degradation. Boxplot showing the distributions of differences in chromatin accessibility frequency (y-axis) upon SOX2 degradation, as a function of the number of SOX2 motifs at CREs (x-axis, color scale). The number of stars represents the significance of the Wilcoxon test performed for each pair of consecutive categories (*ns: p >* 0.05; *\* p ≤* 0.05; *\*\* p ≤* 0.01; *\* *** p ≤* 0.001; *\* * ** p ≤* 0.0001). **(H)** Example locus of an enhancer bound by two copies of SOX2 and one of OCT4. The loss of TF binding at the SOX2 motifs is associated with a reduction in chromatin accessibility frequency in the cell population. The upper panel shows the average SMF signal (1 *−* methylation%) for non-treated and treated samples (black dots and line for untreated, red dots and line for 2h dTAG), as well as TF motifs. The lower panels show *FootprintCharter* stacks for untreated (left) and 2h dTAG (right) samples.

We then excluded CTCF and focused on the remaining 4,535 motifs with differential affinities across alleles, leading to >100 instances of perturbations for most of the 18 tested TFs in our system (Supplementary Figure 3B). We ranked bound TF motifs by their chromatin accessibility frequency on the reference allele and compared it to the frequency of chromatin accessibility on the alternative allele. We observed that most motif disruptions led to a moderate reduction in accessible molecules (<10% median, Supplementary Figure 3M), while complete loss of chromatin accessibility at CREs was rare (Figure 3B). We then stratified CREs based on the number of TFs binding at the locus and quantified how disrupting individual TF motifs affected the width and frequency of chromatin accessibility (Figure 3C). Regardless of the number of TFs binding the CRE on the reference allele, we observed a consistent drop in chromatin accessibility frequency. Upon loss of a single motif, CREs with *n* motifs exhibited distributions of chromatin accessibility frequencies for the alternative allele that approached the distributions of CREs with *n−* 1 motifs on the reference allele. For instance, CREs with 3 motifs had a frequency of accessibility of 50% on the reference allele, and 39% on the alternative allele when one TF motif was disrupted by an SNV (Figure 3C). This frequency was close to that of CREs bound by 2 TFs on the reference allele (35%). These moderate shifts in the frequency of accessible molecules could be observed at individual loci, such as an example of an 11% reduction in chromatin accessibility associated with the loss of a KLF4 binding site (Figure 3D). These observations suggest that loss of individual motifs leads to partial reductions in chromatin accessibility, with each motif contributing cumulatively and rarely more than 10% of the total chromatin accessibility at the locus. To confirm these observations with an orthogonal approach, we quantified the changes in chromatin accessibility frequency at SOX2-bound enhancers upon rapid removal of >90% of SOX2 using a degron system (Figure 3E, Supplementary Figure 3N-O). We observed good agreement in the changes in chromatin accessibility frequency when comparing SOX2 degron data with the SOX2 binding site deletions in our F1 system (Supplementary Figure 3P). Despite the almost complete removal of SOX2, we observed only a partial reduction in the frequency of chromatin accessibility, where enhancer occupancy by nucleosomes rarely increased by more than 10% of the molecules (Figure 3F, Supplementary Figure 3Q). We next asked whether multiple binding sites for the same TF would lead to cumulative effects. When stratifying our data by the number of SOX2 binding motifs present at the CRE, we observed a greater loss of chromatin accessibility at enhancers containing multiple SOX2 binding sites (Figure 3G). This loss can, for instance, be observed at an individual enhancer bound by SOX2 at two distinct motifs where the frequency of molecules with chromatin accessibility decreased by 20% upon SOX2 deletion (Figure 3H). This suggests that the presence of multiple motifs at CREs leads to a cumulative increase in the frequency of chromatin accessibility. Together, these data support the model that the frequency of chromatin accessibility of most CREs is not defined by individual TFs. Instead, the cumulative function of multiple TFs determines the fraction of molecules where chromatin is open for activation.

### Measuring chromatin accessibility of hundreds of enhancers in parallel at an ectopic context

Each TF binding event occurs in a unique genomic environment characterized by specific partner TFs, as well as defined levels of activating and repressing chromatin modifications. This diversity of genomic contexts makes it difficult to tease apart the contribution of the binding of specific TFs on chromatin accessibility from that of the chromatin context or transcriptional activity. Understanding how the chromatin environment impacts TF binding and chromatin accessibility requires a system to study TF function in a defined chromatin context where the influence of the genomic environment is minimized. To that end, we developed a Multiplexed Chromatin-Integrated Reporter Assay (MCHIRA) to measure the ability of hundreds of CREs to establish chromatin accessibility at a defined ectopic genomic locus (Figure 4A). Libraries containing hundreds of CREs were synthesized and cloned in a receiving plasmid flanked by inverted lox sites, which enable its genomic insertion through Recombination Mediated Cassette Exchange (RMCE). The library was inserted at a landing pad located in a neutral chromatin environment that is devoid of either activating or repressive chromatin modifications in mESCs (Supplementary Figure 4A-B). This landing locus was previously used to study the establishment of epigenetic marks, TF binding, and transcription regulation (9, 33–36).

**Fig. 4.**
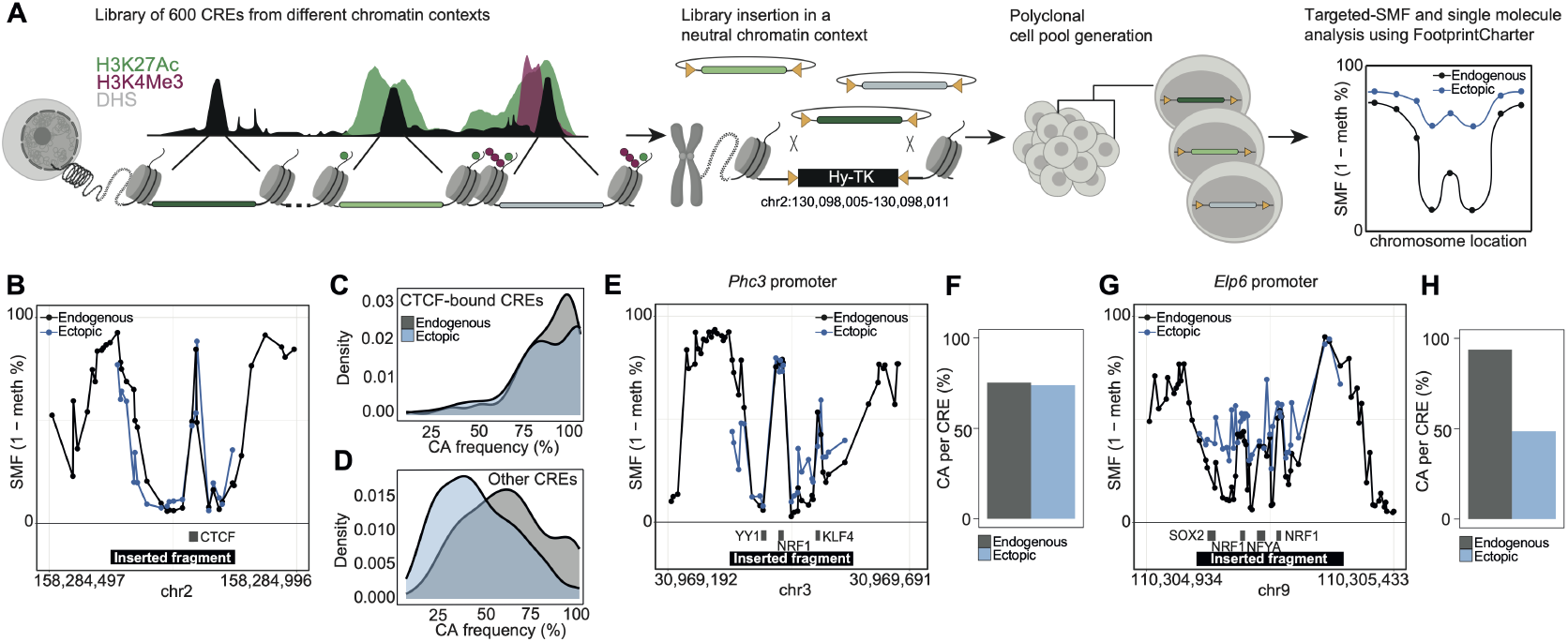
Testing the contribution of the chromatin context to the frequency of chromatin accessibility at CREs. **(A)** Schematic representation of the experimental strategy used to study the effect of the chromatin environment on CRE accessibility. A library of 600 DNA fragments representing entire CREs is synthesized and cloned into a plasmid flanked by asymmetric tandem lox sites for Recombination Mediated Cassette Exchange (RMCE). The fragments are inserted as a pool at the landing site. Negative cell selection enriches for cells that correctly exchanged the Hy-TK (Hygromycin-Thymidine Kinase) cassette by recombination, generating a polyclonal cell population where each cell contains only one insert. The frequency of chromatin accessibility of the inserted fragments is measured by targeted SMF and compared to that measured at the identical genetic sequence at its endogenous locus, as depicted in the cartoon. **(B)** CTCF binding sites are sufficient to establish chromatin accessibility at the ectopic locus. Average SMF signal (1 - methylation %) of a CTCF-containing fragment (black rectangle) at the endogenous (black dots and line) and ectopic locus (blue dots and line). **(C-D)** Chromatin accessibility at the ectopic locus is shifted to lower frequencies, with the exception of CTCF-bound CREs. Density plots of the distribution of chromatin accessibility frequencies at the endogenous (grey) and ectopic (blue) loci for the **(C)** CTCF-bound fragments contained in the library (n = 111) and **(D)** all the non-CTCF-containing fragments (n = 286). **(E-H)** Example loci that **(E)** fully or **(G)** partially establish chromatin accessibility at the ectopic locus. Same representation as (B). For each locus, bar plots show the quantification of chromatin accessibility frequency per CRE at the endogenous (black) and ectopic locus (blue), as measured by *FootprintCharter* **(F)** and **(H)**.

Successful insertion events were enriched by negative selection, avoiding the potential influence of the active transcription of a positive selection marker on the chromatin accessibility of the tested constructs. Chromatin accessibility of the inserted fragments was profiled by targeted SMF using PCR primers that anneal to a synthetic flanking sequence, unambiguously distinguishing the ectopic site from its endogenous counterpart. The frequency of accessibility at the ectopic locus was calculated using *FootprintCharter*, generating single-molecule accessibility data that are directly comparable to our genome-wide SMF measurements. This led to reproducible measures of chromatin accessibility for hundreds of individual fragments of our library in parallel (Supplementary Figure 4C). For each experiment, we profiled chromatin accessibility by SMF at four endogenous loci alongside the ectopic locus. These serve as internal controls to detect potential batch effects in the footprinting procedure and ensure that the ectopic data are comparable across experiments (exemplified in Supplementary Figure 4D).

We designed a library of 600 DNA fragments covering the entire stretch of chromatin-accessible DNA of each CRE (see Methods; 250 bp long). The library was designed to target enhancers and promoters bound by a variety of TFs in diverse chromatin contexts. We obtained sufficient coverage for 66% (397) of the 600 inserted fragments (see Methods and Supplementary Table 2 for library composition). As a positive control, we included CTCF-bound loci, as CTCF motifs can autonomously guide chromatin opening (9). Indeed, we observed high similarity between ectopic and endogenous loci that contained CTCF motifs, with a strong TF footprint and high levels of chromatin accessibility (Figure 4B-C). This demonstrates the ability of our method to accurately quantify the frequency of chromatin accessibility in parallel for multiple sequences inserted at the same ectopic locus.

### Regulatory genetic information only partially determines the frequency of chromatin accessibility

We next systematically compared the frequency of chromatin accessibility of all the non-CTCF-containing ectopic CREs with their genetically identical endogenous counterparts. Overall, 97% of the ectopic sites were accessible above background level, defined as the accessibility observed at unbound motifs (see Methods). Of these, 42% showed comparable frequencies at both loci (Supplementary Figure 4E), while 58% showed less accessibility at the ectopic locus than the endogenous counterpart (Figure 4D, Supplementary Figure 4E). This difference is also evident at individual loci. While chromatin accessibility and TF footprints are recapitulated to near-identical levels at the *Phc3* promoter (Figure 4E-F), they are only partially recapitulated at the *Elp6* promoter (Figure 4G-H). When stratifying by CRE type, we detected a significant reduction of chromatin accessibility at both enhancers and promoters (Supplementary Figure 4F). These results show that the genetic information contained in a 250 bp-long CRE is generally sufficient to partially open chromatin, yet for the majority of CREs, the frequency of accessibility is lower than at the endogenous site. To understand if the presence of specific TF motifs determines the autonomy of certain sequences to recapitulate chromatin accessibility at the ectopic locus, we compared the accessibility levels of CREs at endogenous and ectopic sites, stratifying based on the presence of specific TF motifs within the fragment (Figure 5A, Supplementary Figure 4G). With the exception of CTCF, we observed that regardless of the TF motifs contained in the fragment, the frequency of accessibility is lower at the ectopic versus the endogenous loci (Figure 5A). For example, fragments containing NFY and NRF1 motifs reached 68% and 60% accessibility on average at the ectopic site compared to >80% at endogenous loci (Figure 5A, Supplementary Figure 4G). Similarly, fragments containing other TF motifs, such as SOX2 or KLF4, show a reduction in accessibility frequency when inserted at the ectopic site (Figure 5A). In conclusion, with the notable exception of CTCF, the presence of most TF motifs is not sufficient to recapitulate endogenous chromatin accessibility. This includes TFs associated with higher frequencies of chromatin accessibility at endogenous loci, such as NFY and NRF1.

**Fig. 5.**
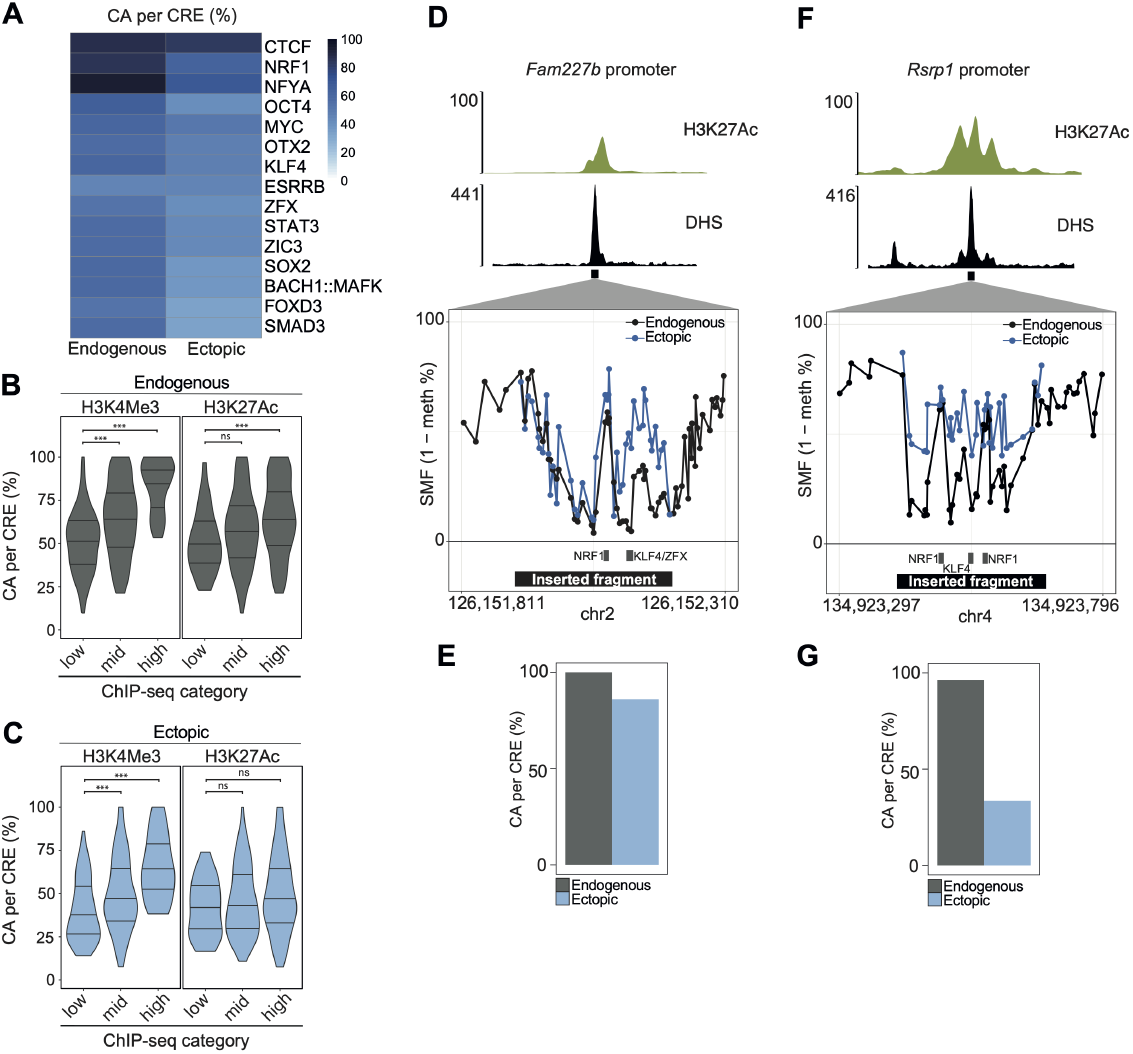
Identification of the genetic and epigenetic determinants of chromatin accessibility. **(A)** TFs bind to their motifs and open chromatin at the ectopic locus, but generally to a lower extent compared to the endogenous motifs. Heatmap showing the median values of chromatin accessibility (CA) at the endogenous and ectopic loci as a function of the presence of specific TF motifs within the DNA fragments, as defined by ChIP-seq and motif annotation (see Methods). **(B-C)** The frequency of chromatin accessibility (CA) scales with chromatin modification levels at the endogenous loci, but not at the ectopic site. **(B)** Violin plots showing the frequency of chromatin accessibility at CREs when measured at the endogenous loci (black) as a function of H3K4Me3 (left panel) or H3K27Ac (right panel) ChIP-seq enrichment as measured at the endogenous loci (*n* = 286; ns: *p >* 0.05; *: *p ≤* 0.05; **: *p ≤* 0.01; ***: *p ≤* 0.001; ****: *p ≤* 0.0001). **(C)** Similar to (B), but depicting the frequency of chromatin accessibility (CA) at CREs when measured at the ectopic locus (blue). **(D)** Single example of the *Fam227b* promoter that faithfully recapitulates chromatin accessibility at the ectopic locus. The upper panel shows genome browser tracks of two promoter regions that are characterized by low (left) and high (right) levels of H3K4Me3 and H3K27Ac endogenously at the DHS peak. The lower panel shows the average SMF signals showing the endogenous and ectopic chromatin accessibility frequencies (ns: *p >* 0.05; *: *p ≤* 0.05; **: *p ≤* 0.01; ***: *p ≤* 0.001; ****: *p ≤* 0.0001). **(E)** Quantification of the frequency of chromatin accessibility for the same fragment at the endogenous (black) and ectopic locus (blue), as measured by *FootprintCharter*. **(F-G)** Single example of the *Rsp1* promoter that partially recapitulates chromatin accessibility at the ectopic locus. Same representation as (D-E).

Together, our data suggest that frequencies of chromatin accessibility are partially encoded in the genetic sequence through TF motifs, but that additional mechanisms may exist at endogenous loci to promote higher accessibility frequencies of CREs.

### Deposition of H3K27Ac increases the frequency of chromatin accessibility

Since the regulatory sequences inserted at the ectopic locus are genetically identical to their endogenous counterparts, we hypothesized that the observed reduced accessibility at the ectopic site could be explained by differences in the chromatin context. While most CREs are marked with the active chromatin modifications H3K4me3 and/or H3K27Ac at their endogenous location, these marks are absent at the ectopic insertion site (Supplementary Figure 4B). When we stratified the endogenous CREs based on their H3K4me3 and H3K27Ac ChIP-seq signal, we observed a scaling between the level of these two active marks and the frequency of chromatin accessibility (Figure 5B; Supplementary Figure 4H-I). In contrast, we observed this correlation only for H3K4me3 (Figure 5C, left panel) but not for H3K27Ac (Figure 5C, right panel) when comparing the presence of the mark at the endogenous locus with the frequency of chromatin accessibility at the ectopic locus.

This difference is particularly evident when comparing two NRF1-bound promoters with different endogenous levels of H3K27Ac (Figure 5D and Figure 5F). The promoter from the *Fam277b* locus, which has low H3K27Ac, almost perfectly recapitulates chromatin accessibility at the ectopic site (Figure 5D-F), while the promoter fragment from the highly acetylated *Rsrp1* locus shows only partial recapitulation (Figure 5F-G). These results suggest a model in which H3K27Ac enhances the frequency of chromatin accessibility at the endogenous locus, an effect that is entirely or partially lost when inserting the sequence at the ectopic locus, which lacks histone modifications.

To test this model, we globally reduced H3K27Ac levels using chemical inhibition of the histone acetyltransferase p300 (A-485 treatment) (37), which binds a subset of enhancers in mESCs (38). We measured changes in H3K27Ac by ChIP-seq and identified 5,823 CREs with significant loss of acetylation after 24 hours of treatment (Supplementary Figure 5A-B). We then quantified the changes in the frequency of chromatin accessibility at these CREs using SMF under the same conditions (Supplementary Figure 5C). We observed that the magnitude of chromatin accessibility loss scales with the reduction of H3K27Ac levels (Figure 6A, Supplementary Figure 5D), suggesting that H3K27Ac may increase chromatin accessibility frequency at these loci. Another prediction of our model is that reducing H3K27Ac deposition should selectively affect chromatin accessibility frequency at endogenous loci, whereas their ectopic counterparts, which are located in a chromatin-neutral environment, should remain unchanged. To test this, we compared the accessibility changes of the p300-sensitive CREs with the response of their matching ectopic CRE insertions from our library. We repeated the p300 inhibition experiment on cell lines containing our CRE libraries and measured changes in chromatin accessibility frequency using targeted SMF (Supplementary Figure 4D, Supplementary Figure 5E). We found that CREs insensitive to p300 inhibition, including CTCF-bound loci, maintain consistent accessibility frequencies across conditions regardless of their genomic location (Figure 6B-C). For example, the *Mctp2* promoter, which has low endogenous levels of H3K27Ac, is unaffected by p300 inhibition at either its endogenous or ectopic location (Figure 6D). In contrast, CREs that respond to p300 inhibition at their endogenous loci no longer lose chromatin accessibility upon p300 inhibition when inserted at the ectopic site (Figure 6C). For instance, the *Xpa* enhancer, which has high endogenous levels of H3K27Ac, loses approximately 20% of its accessibility upon p300 inhibition at the endogenous locus but remains unchanged at the ectopic site (Figure 6E). These findings confirm that H3K27Ac enhances chromatin accessibility frequency at endogenous CREs, while ectopic inserts primarily rely on TF binding to open chromatin. Together, our data suggest that the frequency of chromatin accessibility at CREs is defined by the cumulative action of multiple TFs and is further enhanced by the deposition of histone marks such as H3K27Ac.

**Fig. 6.**
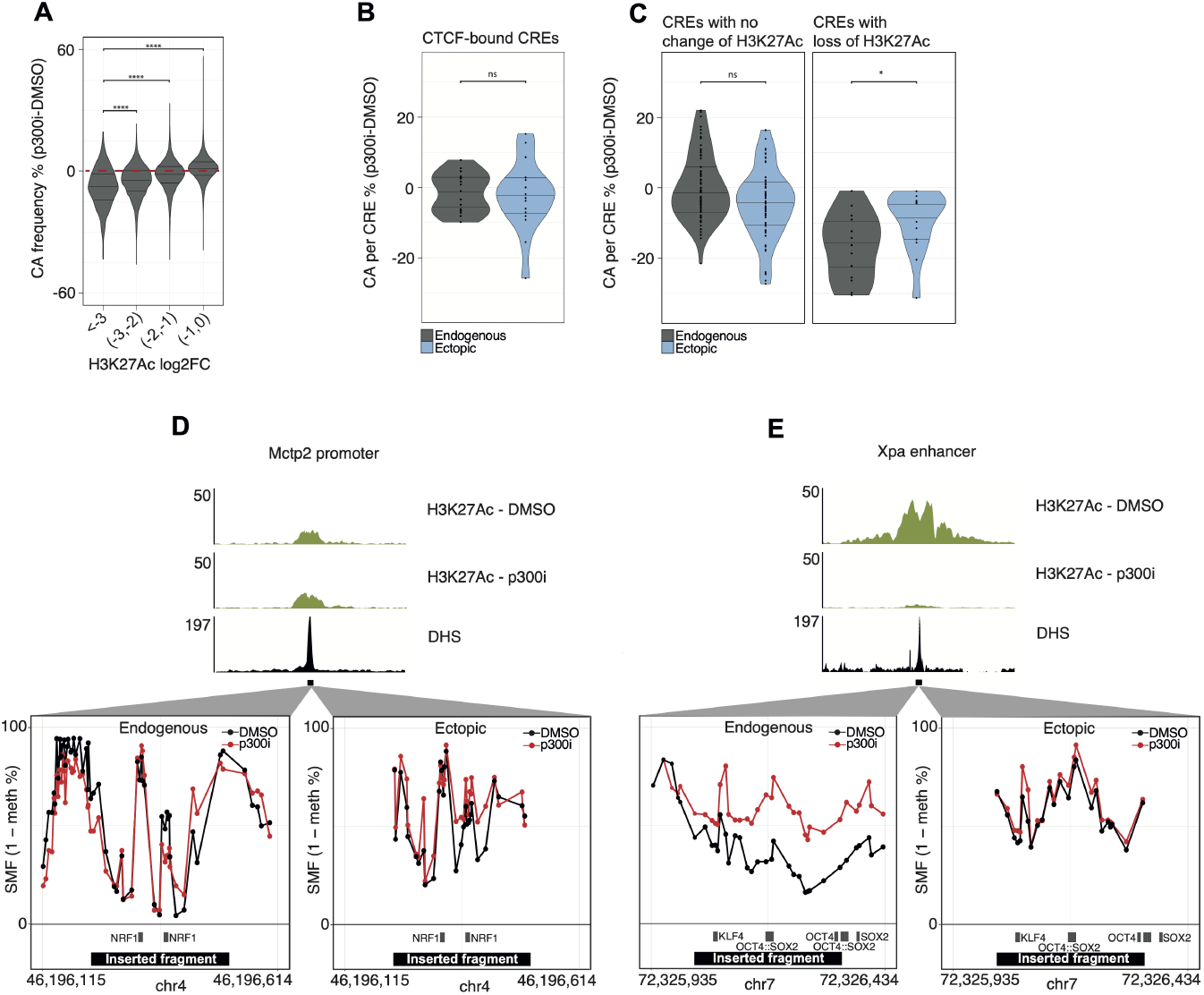
H3K27Ac enhances the frequency of chromatin accessibility at enhancers. **(A)** Loss of chromatin accessibility scales with the loss of H3K27Ac at endogenous loci. Violin boxplots showing the difference in the frequency of chromatin accessibility (CA_p300i_ *™* CA_DMSO_) at endogenous loci as a function of H3K27Ac log_2_ fold changes (*n* = 73, 529; ****: *p ≤* 0.0001). **(B)** CTCF is not affected by p300 inhibition. Violin boxplots showing the difference in the frequency of chromatin accessibility (CA_p300i_ *™* CA_DMSO_) at the endogenous (black) and ectopic loci (blue) upon p300 inhibition (Statistical testing was performed using a one-sided Wilcoxon test; ns: *p >* 0.05; *: *p ≤* 0.05; **: *p ≤* 0.01; ***: *p ≤* 0.001; ****: *p ≤* 0.0001). **(C)** p300 inhibition affects the frequency of chromatin accessibility at endogenous CREs, but not at the ectopic locus. CREs were categorized based on the significant change in H3K27Ac signal at the endogenous loci upon p300 inhibition. Similar representation as (B). **(D)** Single locus example of the changes in chromatin accessibility upon p300 inhibition at a promoter with no H3K27Ac loss when measured at the endogenous (left) or ectopic (right) locus. The upper panel shows genome browser tracks of H3K27Ac in DMSO and p300 inhibition (p300i) conditions, and DHS. The lower panel shows average SMF signals (1-methylation %) in control (DMSO – black line and dots) and cells treated with a p300 inhibitor (p300i - red dots and line). **(E)** Single locus example of the changes in chromatin accessibility upon p300 inhibition at an enhancer regulated by p300 when measured at the endogenous (left) or ectopic (right) locus. Same representation as (D).

### Frequency of chromatin accessibility defines Pol II recruitment levels at enhancers

Chromatin accessibility is an indirect proxy for enhancer activity, as many open chromatin regions are not transcriptionally active. We next examined how differences in chromatin accessibility frequency at enhancers relate to their transcriptional activation potential. As a proxy for enhancer activity, we measured the recruitment of transcriptionally active RNA polymerase II (Pol II) at enhancers using precision nuclear run-on sequencing (qPRO-seq). While indirect, measuring active Pol II circumvents the challenges associated with assigning enhancers to target genes when profiling transcriptional changes (27, 39). We performed allele-resolved qPRO-seq in both F1 lines and quantified Pol II recruitment changes at CREs. We then compared these changes to differences in chromatin accessibility frequency observed between alleles in response to TF motif deletions. We observed that decreases in chromatin accessibility frequency led to reduced levels of active Pol II at enhancers and promoters (Figure 7A). At enhancers, this decrease in activity scaled with the magnitude of accessibility frequency changes (Figure 7A). Furthermore, even modest differences in chromatin accessibility frequency between alleles, such as those caused by the loss of individual TFs, were associated with significant changes in enhancer activity (Figure 7A, exemplified in Figure 7B). These findings demonstrate that the frequency of chromatin accessibility at enhancers determines the frequency at which they are transcriptionally active through the recruitment of Pol II.

**Fig. 7.**
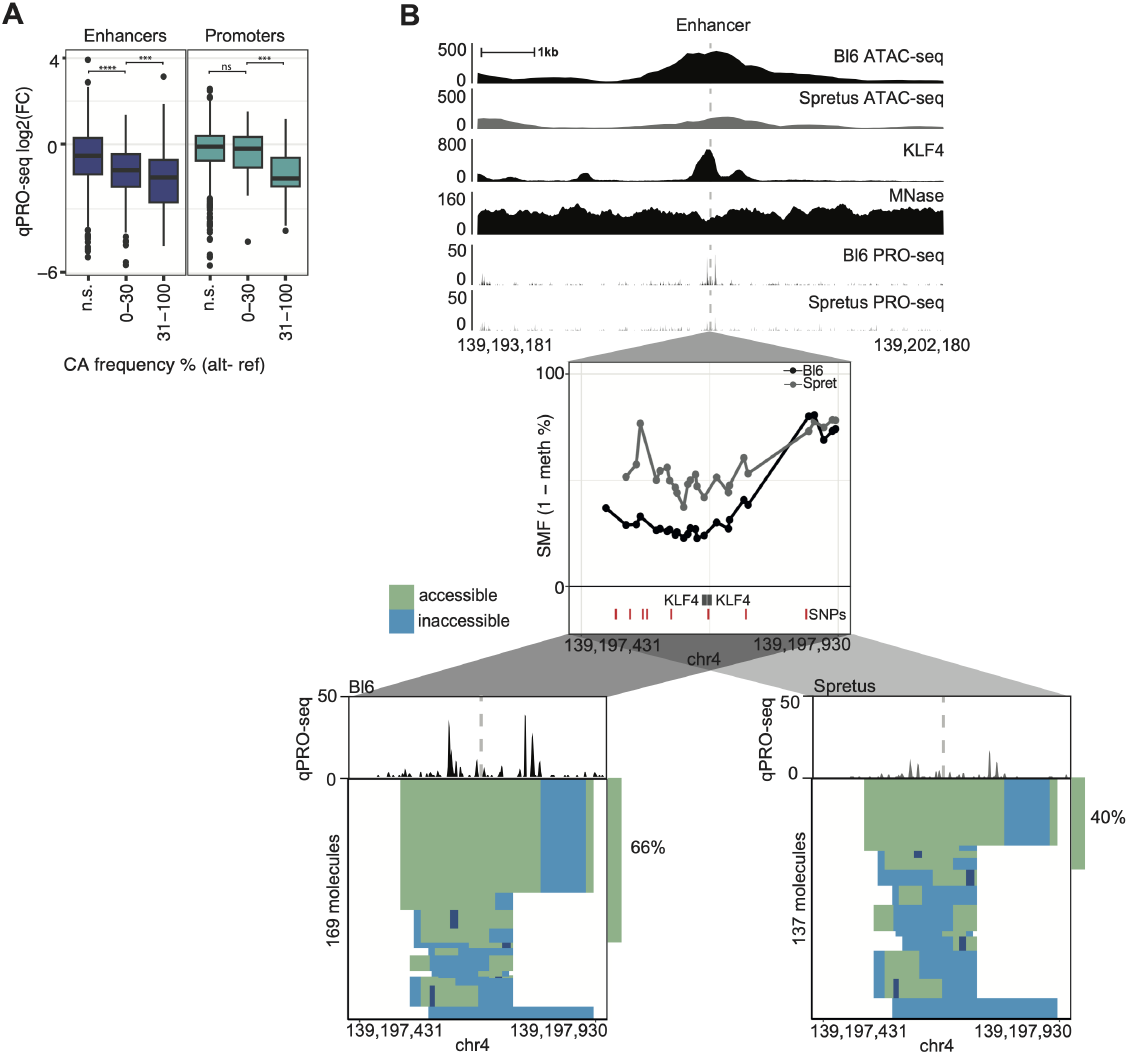
Pol II activity of enhancers depends on the frequency of chromatin accessibility. **(A)** Boxplots showing the distributions of qPRO-seq log_2_ fold changes as a function of chromatin accessibility frequency reductions for TF motifs with allele-specific affinities at enhancers (left panel) or promoters (right panel). The number of stars represents the significance of the Wilcoxon test performed for each pair of consecutive categories (ns: *p >* 0.05; *: *p ≤* 0.05; **: *p ≤* 0.01; ***: *p* 0.001; ****: *p≤* 0.0001). **(B)** Example enhancer bound by two active KLF4 motifs, one of which has a strong difference in affinity between the Bl6 and Spret alleles. This genetic variation is associated with a reduction in chromatin accessibility frequency of 26% in the cell population, leading to a reduction in qPRO-seq signal at the enhancer. The upper panel shows genomic tracks for BL6xSpret F1 mESCs ATAC-seq (Bl6 in black, Spretus in grey), mESCs ChIP-nexus for KLF4 and MNase-seq, and BL6xSpret F1 mESCs qPRO-seq (Bl6 in black, Spretus in grey) around the TF motifs. The middle panel shows the average SMF signal (1-methylation %) for the BL6xSpret F1 line (Bl6 in black, Spretus in grey), as well as the TF motif and SNVs annotation. The lower panels show *FootprintCharter* stacks for the Bl6 (left) and Spretus (right) alleles, along with qPRO-seq signal at the locus.

## Discussion

In this study, we leveraged SMF to dissect the mechanisms regulating the frequency at which chromatin is accessible at enhancers in a cell population. Our findings reveal that chromatin accessibility at enhancers results from the cumulative action of multiple transcription factors (TFs) rather than the function of individual specialized TFs. By measuring the ability of regulatory genetic elements to establish chromatin accessibility outside of their endogenous genomic context, we demonstrate that chromatin accessibility at CREs is only partially encoded in the genetic sequence. Activating chromatin modifications, such as H3K27Ac, further enhance the frequency at which chromatin is open. Moreover, we establish that chromatin accessibility frequency determines the levels of RNA polymerase II (Pol II) recruitment at enhancers, thereby regulating their potential to activate target genes. We observed that depletion of most individual TFs leads to only a partial loss of chromatin accessibility, consistent with previous reports (32). Additionally, the requirement for cumulative TF binding to maintain chromatin accessibility aligns with the observation that many TF motifs cannot independently open chromatin (9). Notably, our data show that individual TFs promote chromatin accessibility in approximately 10% of cells on average, concordant with recent findings using synthetic constructs containing increasing numbers of KLF1 motifs (40). These findings support a ‘cumulative model’, wherein multiple TFs act additively to continuously maintain chromatin accessibility. This model complements the ‘pioneer’ model, where a subset of TFs initially engage with nucleosomes to open chromatin. In this sequential framework, broader TF participation in maintaining an open chromatin state may be facilitated by the presence of destabilized nucleosomes (41), which remain indistinguishable from canonical nucleosomes in our data. SMF provides a snapshot of DNA occupancy in a cell population, where the frequency of an event is determined by both its occurrence rate and duration on DNA (38). TF residence time is typically within seconds, yet ATP-dependent chromatin remodeling complexes must be continuously recruited to maintain open chromatin, as nucleosomes tend to reoccupy sites within minutes upon inhibition of remodeling activity (12, 13). Increasing the number of TF binding sites within an enhancer may extend the duration of TF occupancy at CREs, thereby sustaining chromatin accessibility. This cumulative function of transient TF occupancy events could prolong chromatin accessibility over extended periods, similar to the TF exchange model proposed to explain how transcriptional bursts exceed TF residence time on DNA (42). Studying regulatory genetic elements outside their native genomic context revealed that enhancer activity frequency is genetically encoded via TF motifs, yet the observed chromatin accessibility levels in the genome are further enhanced by activation marks such as H3K27Ac. This suggests a model in which chromatin modifications are not strictly required for chromatin opening but instead enhance the frequency of enhancer accessibility and Pol II recruitment. Our data identify a large set of short CREs capable of recruiting TFs but insufficient for efficient p300 recruitment and H3K27Ac deposition. Dissection of p300’s protein interaction domains suggests that efficient recruitment requires interactions with multiple TFs (43). It is possible that effective p300 recruitment necessitates more than the 2–4 TF binding sites typically present in our CREs, or even cooperative interactions between adjacent CREs.

Our findings that H3K27Ac enhances chromatin accessibility align with reports indicating that H3K27Ac reduces the TF concentration required for SOX9 function (19). These data provide a potential molecular explanation for the effects of chromatin environment on enhancer activity. Modulation of accessibility frequency by activating chromatin marks could account for enhancer activity shifts observed at various loci in multiplexed reporter assays (44–46). However, the mechanisms by which acetylation enhances chromatin accessibility remain to be elucidated. Potential explanations include increased TF binding affinity (47) or recruitment of bromodomain-containing chromatin-modifying enzymes (48).

## Methods

### Cell Lines

159 WT mESCs (49) and 159 DNA methyltransferase triple-knockout (DNMT TKO, (50) were used to generate the reference SMF datasets (15). Bl6xSpretus F1 hybrid cells were obtained from (51). F1 hybrid cells Bl6xCast were derived at the DKFZ mouse facility (Odom lab). TC-1 mESC containing the landing pad for RMCE (background 129S6/SvEvTac, originally obtained from A. Dean at the National Institutes of Health). All three cell lines were targeted for CRISPR-KO of the three DNA methyl transferases to generate Bl6xCast DNMT TKO and Bl6xSpret DNMT TKO and TC-1 TKO. SOX2 degron mESC line were previously derived (52). See Supplementary Table 3 for a list of the experiments performed in this study and the corresponding cell line.

### Generation of DNMT Knockout Lines

DNMT1/3a/3b TKO cells were generated in TC-1 and F1 hybrid (Bl6xCast and Bl6xSpretus) cell lines using gRNAs previously described (50). The three gRNAs targeting each DNMT gene were cloned into pX459 (Addgene plasmid #48139) and transfected into the TC-1 and F1 hybrid (Bl6xCast and Bl6xSpretus) cell lines. After puromycin selection, the pool was analyzed by PCR-RFLP (Restriction Fragment Length Polymorphism) and HpyCH4V digest to confirm DNMT3a/b double knockout. Subsequently, 96 colonies were picked and screened individually by PCR-RFLP and HpyCH4V digest. DNA triple mutant clones were further confirmed by amplicon sequencing using primer sets specific for DNMT targets. For the TKO clone used in this study, complete loss of DNA methylation was confirmed by DNA digestion with the DNA methylation-sensitive restriction enzyme HpaII. Primer Sequences for DNMT Targets: 1) DNMT1: GCAGGAGGGACACAGTCATT, AGGACTGCAACGTGCTTCTT (Tm: 63°C, Amplicon Length: 359 bp); 2) DNMT3a: TTGGCACCTCCCAGGGTTC, TCGGACAGTGAGTGGTGAGG (Tm: 63.6°C, Amplicon Length: 400 bp); 3) DMNT3b: CCCCCAGAGTCCAGTTCTTA, GGAAGGACCGGGCCTGAG (Tm: 64°C, Amplicon Length: 281 bp)

### Single Molecule Footprinting (SMF)

SMF was performed as previously described (15, 29). Briefly, cultured cells were harvested using trypsin and washed twice with 1x PBS. For each reaction, 250,000 cells were used. Cell pellets were resuspended in ice-cold lysis buffer and incubated on ice for 10 minutes before being spun at 1,000x g for 5 minutes at 4°C. Nuclei were resuspended in 1x M.GpC buffer (NEB, #M0227L). For GpC methyltransferase treatment, freshly prepared GpC methyltransferase mix (1x M.GpC buffer, 300 mM sucrose, 64 µM SAM (NEB, #B9003S), M.CviPI (NEB,#M0227L)) was added and incubated at 37°C for 7.5 minutes. After adding the stop solution and proteinase K, the incubation continued overnight at 55°C. DNA extraction was performed the next day using phenol-chloroform, followed by RNase A treatment.

For targeted SMF at the ectopic site, DNA was bisulfite converted using the Qiagen Epitect bisulfite kit as described in (29), and then amplified using KAPA HiFi Uracil+ (Roche). The sequencing library was prepared using the NEBNext DNA Ultra II library preparation kit and sequenced with a MiSeq 250bp paired-end run.

### Western Blotting

Western blotting was performed as follows: cells were harvested, washed in ice-cold PBS, and resuspended in cold Lysis Buffer A (10 mM HEPES, 5mM MgCl2, 0.25M Sucrose, 0.1% NP40). Cytoplasmic membranes were lysed by incubation on ice for 20 minutes, followed by multiple pipetting steps. Nuclei were pelleted at 9800x g for 10 minutes at 4°C, and lysed further in Lysis Buffer B (2.5 mM HEPES, 1.5 mM MgCl2, 0.1 mM EDTA, 20% Glycerol) for 10 minutes. Nuclear extracts were recovered by vortexing for 20 seconds at max power, followed by centrifugation at 15,000x g for 15 minutes at 4°C. Protein concentration was determined using the Qubit protein quantification kit.

For SDS-PAGE, 20µg of protein was loaded onto a 10% gel, transferred to a nitrocellulose membrane, and incubated overnight at 4°C with antibodies against HA (1:1000, Biolegend, #901501) for SOX2 and H3 (1:2000, Cell signaling, #4499T) as a loading control.

### Cross-Link ChIP-seq

The protocol was adapted from (53), with minor modifications. Ten million mESCs were harvested and cross-linked in 3 mL pre-tempered (25°C) ES medium containing 1% formaldehyde and supplemented with sodium butyrate (5 mM final concentration) for 10 minutes at room temperature with rotation. The cross-linking reaction was quenched by adding glycine (125 mM final concentration) and incubated for 10 minutes at room temperature. Cells were washed twice with ice-cold PBS containing 10% FBS and centrifuged at 200x g for 5 minutes at 4°C. At this point, cell pellets can be snap-frozen and stored at −80°C for several months.

Pellets were resuspended in 300 µL Sonication Buffer (50 mM Tris-HCl pH 8.0; 0.5% SDS) and chromatin was sheared using the Bioruptor Pico (Diagenode) for 15 cycles (30” ON/30” OFF). Sonicated lysates were diluted 1:6 in lysis buffer (10 mM Tris-HCl, pH 8.0; 100 mM NaCl; 1% Triton X-100; 1 mM EDTA; 0.5 mM EGTA; 0.1% sodium deoxycholate; and 0.5% N-lauroylsarcosine). After centrifugation at full speed for 10 minutes at 4°C, the soluble fraction was collected. A 2.5% portion of the supernatant was kept as input, and the fragmentation pattern (150-500 bp) was analyzed by agarose gel electrophoresis. At this point, sonicated lysates may be aliquoted and stored at −80°C with a final concentration of 10% glycerol.

For each immunoprecipitation (IP), 30 µL of Protein G Dynabeads (Invitrogen) were washed twice with 1 mL of PBS-T (PBS + 0.01% Tween-20) and incubated with the antibody against H3K27ac (ab4729, Abcam) for 1 hour at room temperature with rotation. Coated beads were washed once in cold PBS-T and twice in lysis buffer, then resuspended in 30 µL of lysis buffer per IP and added to 15 µg of chromatin. At this point, 1 µg of exogenous chromatin (from Drosophila Schneider 2 (S2) cells, prepared with the same protocol and stored at −80°C with a final concentration of 10% glycerol) was added to each reaction as spike-in.

After overnight incubation at 4°C with rotation, bead-immunocomplexes were washed twice (5 minutes per wash), using the following buffers: RIPA, RIPA supplemented with 360 mM NaCl, and LiCl buffer (10 mM Tris-HCl, pH 8.0; 250 mM LiCl; 0.5% NP-40; 0.5% sodium deoxycholate; 1 mM EDTA). The complexes were then briefly rinsed with TE buffer and eluted in ChIP SDS elution buffer (10 mM Tris-HCl, pH 8.0; 300 mM NaCl; 5 mM EDTA; 0.5% SDS).

RNA and protein digestion were performed by adding 2 µL of RNase A (10 mg/mL stock) and incubating for 30 minutes at 37°C, followed by the addition of 1.5 µL of Proteinase K (20 mg/mL stock) and incubation at 55°C for 1 hour. Cross-links were reversed by overnight incubation at 65°C. DNA was then purified using 1.4X SPRI-select beads.

Sequencing libraries were prepared using the NEBNext Ultra II DNA Library Preparation Kit (New England Biolabs) and sequenced on Aviti Cloudbreak Low 2×150 (250 M clusters/run).

### PRO-seq

Precision run-on sequencing (PRO-seq) was performed as previously described (39, 54) with slight modifications. Five million TKO F1 mESCs were used per sample with a spike-in of 0.05 million (1%) Drosophila S2 cells. Two replicates per cell line were generated, with each replicate prepared simultaneously for all cell lines and conditions. Cells were harvested using Trypsin-EDTA on ice and washed twice with 10 mL ice-cold PBS, followed by centrifugation at 1000x g for 4 minutes at 4°C. Permeabilization was performed in 10 mL of permeabilization buffer (10 mM Tris-HCl pH 8.0, 250 mM sucrose, 10 mM KCl, 5 mM MgCl_2_, 1 mM EGTA, 0.1% Igepal, 0.05% Tween-20, 0.5 mM DTT, 10% (vol/vol) glycerol, 1 tablet of protease inhibitor per 50 mL, 4 units/mL SUPERaseIN inhibitor) on ice for 10 minutes. The cell pellets were then washed twice with 10 mL of ice-cold cell wash buffer (10 mM Tris-HCl pH 8.0, 250 mM sucrose, 10 mM KCl, 5 mM MgCl_2_, 1 mM EGTA, 0.5 mM DTT, 10% (vol/vol) glycerol, 1 tablet of protease inhibitor per 50 mL, 4 units/mL SUPERaseIN inhibitor) with centrifugation at 1000x g for 4 minutes at 4°C. Cells were resuspended in 50 µL freeze buffer (50 mM Tris-HCl, pH 8.0, 40% (vol/vol) glycerol, 5 mM MgCl_2_, 1.1 mM EDTA, 0.5 mM DTT and 4 units/mL SUPERaseIN inhibitor), snapfrozen, and stored at −80°C until further use.

The nuclear run-on reaction was performed as a 2-biotin run-on. For this, 50 µL of the 2x Nuclear-run on master mix (10 mM Tris-Cl pH 8.0, 5 mM MgCl_2_, 1 mM DTT, 300 mM KCl, 40 µM Biotin-11-UTP, 40 µM Biotin-11-CTP, 40 µM ATP, 40 µM GTP, 1% Sarkosyl, 1 µL SUPERaseIN inhibitor) was prepared for each sample. The reaction mix was preheated at 37°C and 50 µL of the pre-calculated number of cells were added to each reaction vial, mixed thoroughly and incubated at 37°C for 5 minutes with shaking (750 rpm). To stop the reaction, 350 µL of the RL Buffer from the Norgen RNA extraction kit were added and vortexed. 240 µL of 100% ethanol was added to the mixture and vortexed again. RNA extraction was performed according to the kit’s manual. The final RNA was eluted twice with 50 µL H_2_O and pooled to a final volume of 100 µL.

For the base hydrolysis, the RNA was denatured for 30 seconds at 65°C and snap-cooled on ice. 25 µL of ice-cold 1N NaOH were added and incubated on ice for 10 minutes. RNA was precipitated by adding 125 µL Tris-HCl (pH 6.8), 5 µL NaCl, 1 µL GlycoBlue and 650 µL 100% EtOH and centrifugation at 20,000x g at 4°C for 20 minutes. The RNA pellet was washed with 70% EtOH, air-dried and resuspended in 6 µL H_2_O.

For the 3’ RNA adaptor ligation, 1 µL of REV3 3’RNA adaptor dilution (10 µM), heat-denatured at 65°C for 20 seconds and placed on ice, was added to the 6 µL resuspended RNA. 13 µL of the RNA ligation mix using T4 RNA ligase I was added and incubated at 25°C for 1 hour. Biotin RNA enrichment was performed by adding 55 µL binding buffer (10 mM Tris-HCL pH 7.4, 300 mM NaCl, 0.1% Tergitol, 1 mM EDTA) to each ligation reaction, followed by 25 µL of pre-washed streptavidin beads. Reactions were incubated on a rotator set to 8 rpm at room temperature for 20 minutes. Beads were washed once with ice-cold high-salt buffer (50 mM Tris-HCL pH 7.4, 2M NaCl, 0.5% Tergitol, 1mM EDTA), transferred to new tubes, and washed once with low-salt buffer (5 mM Tris-HCL pH 7.4, 0.1% Tergitol, 1mM EDTA).

### Library Design and Cloning

To construct the DNA libraries, we selected 600 candidate cis-regulatory elements (CREs) bound by transcription factors (TFs) of interest, including pluripotency and ubiquitous TFs. Binding of these TFs to the selected CREs was confirmed using published ChIP-sequencing data (see Supplementary Table 1 for TFs of interest and ChIP-seq datasets, and see Supplementary Table 2 for library composition).

Additionally, the chosen CREs were required to contain a sufficient density of CpG and GpC dinucleotides to ensure adequate single-molecule footprinting (SMF) resolution, while avoiding an overrepresentation of CpG islands. The selected 250 bp DNA fragments were synthesized as ssDNA oligo pool by Twist Bioscience. The oligo pool was PCR amplified following the manufacturer’s instructions. The library was cloned using In-Fusion seamless cloning (Takara Bioscience) into a receiving plasmid containing asymmetric tandem Lox sites, for targeted insertion into the genome through Recombination-Mediated Cassette Exchange (RMCE), and priming regions for targeted SMF.

### Library Insertion

We used previously described TC-1 ES cells that have a landing pad for targeted insertion using RMCE (36). We used targeted locus amplification coupled with next-generation sequencing (Cergentis) to genotype the precise location of the landing pad. We mapped the landing pad at mouse chr2:130,098,005-130,098,011 (Supplementary Figure 4A and Supplementary Figure 4B) within a gene-poor region with no active transcription, no active chromatin marks, and no repressive chromatin marks.

For targeted insertion of the DNA libraries into the mouse genome, RMCE was used as previously described (35). Briefly, TC-1 DNMT TKO ES cells were selected under hygromycin (250 µg/mL, Roche, Switzerland) for 14 days. Afterward, 50 million cells were nucleofected (Amaxa 4D-Nucleofector Core Unit, Lonza) with 300 µg DNA libraries containing plasmid and 180 µg pIC-CRE plasmid.

After 2 days, cells were selected with 3 µM ganciclovir (Selleckchem, Catalog No. S1878) for 10 days.

### PRO-seq Data Pre-processing

For the F1 PRO-seq data generated in this study, raw sequencing reads were trimmed using Cutadapt (version 3.5) (55). UMIs were detected and extracted using UMI-tools (version 1.1.2) (56).

To assign pre-processed reads to the correct allele of origin, SNVs annotation between the species of interest were fetched from the Mouse Genome Project portal (57). SNVs were injected into the UCSC *Mus musculus* reference genome distributed through the Bioconductor (58, 59) package BSgenome.Mmusculus.UCSC.mm10 (60) using the function *qAlign* provided with the Bioconductor package QuasR (61). This resulted in two separate genomes to which reads were competitively aligned using the QuasR function *qAlign* with default alignment parameters (61).

Aligned reads were deduplicated using UMI-tools. Reads aligned to rRNAs were removed.

### Processing of Publicly Available ChIP-seq and ChIP-nexus Datasets

Publicly available ChIP-seq and ChIP-nexus datasets were pre-processed using the nf-core pipeline nf-core/chipseq v.2.0.0 with arguments “– narrow_peak –read_length 50” (62). For our H3K27Ac and H3K4Me3 ChIP-seq analysis, we calculated the coverage within 2kb windows using the *qCount* function of the QuasR package, focusing on Transcription Factor Binding Sites (TFBS) in genomic regions enriched during the SMF bait capture step. Based on the enrichment patterns of H3K27Ac and H3K4Me3, we categorized the regions into three classes: low, mid, and high enrichment. For H3K27Ac, classes are as follows: low (<100), mid (between 100 and 500), and high (>500). For H3K4Me3, classes are as follows: low (<50), mid (between 50 and 300), and high (>300). See Supplementary Figures 4H-I.

### ChIP-seq upon p300 Inhibition Pre-processing and Analysis

Illumina adaptors and low-quality bases were trimmed using TrimGalore (v 0.6.7). Pre-processed reads were aligned to the mouse reference genome (BSgenome.Mmusculus.UCSC.mm10) using the R package QuasR, which uses Bowtie as the aligner (alignment parameter: -m 1 -e 70 -X 1000 -k 2 –best –strata). Only uniquely aligned reads were kept and deduplicated using the tool *MarkDuplicates* from Picard (v 3.1.0). Normalized signal coverage bigwig files were obtained with deepTools *bamCoverage* (v 3.5.0) and a bin size of 50 bases.

The H3K27Ac ChIP-seq contained an exogenous spike-in, for which reads were aligned independently to the Drosophila reference genome (BSgenome.Dmelanogaster.UCSC.dm6) and processed as described for the reads aligned to the mouse genome. The number of uniquely mapped reads aligning to major mouse and fly chromosomes, in both IP and input samples, was obtained and used to compute normalization factors (using the formula: aligned reads * (target IP/spike-in IP) * (spike-in input/target input) (63)). The normalization factors were then applied with deepTools *bamCoverage* to generate scaled bigWig files.

The log2 fold change of H3K27Ac and statistical testing were computed using the DESeq2 Bioconductor package (64), providing H3K27Ac counts (calculated using the *qCount* function of the QuasR package, in 2kb windows covering the genomic regions enriched during the SMF bait capture step) in DMSO or p300i treatment replicates, after applying the normalization factors. The p-values were adjusted using the Benjamini-Hochberg method.

### F1 Bl6xCast ATAC-seq Data Pre-processing

Publicly available ATAC-seq data produced using an analogous Bl6xCast F1 ESC line (65) was fetched using the nf-core pipeline nf-core/fetchngs (66). Sequencing reads were trimmed for Illumina adapters and low-quality bases using TrimGalore! v0.6.7 (67). Reads shorter than 20bp were discarded after trimming.

To assign pre-processed reads to the correct allele of origin, SNVs annotations between mouse species were fetched from the Mouse Genome Project portal (57). SNVs were injected into the UCSC *Mus musculus* reference genome distributed through the Bioconductor (58, 59) package BSgenome.Mmusculus.UCSC.mm10 (60) using the function *qAlign* provided with the Bioconductor package QuasR (61). This resulted in two separate genomes to which reads were competitively aligned using the QuasR function *qAlign* with alignment parameters “-e 70 -X 1000 -k 2 –best –strata”. The best alignment determined the allelic assignment for each read. Finally, duplicated reads were identified and discarded using the Picard tool *MarkDuplicates* v2.15.0 (68).

### Bait Capture SMF Data Pre-processing

Bait capture SMF sequencing reads were trimmed for Illumina adapters and low-quality bases using TrimGalore v0.6.7. Reads shorter than 20bp after trimming were discarded.

For F1 SMF data, to assign pre-processed reads to the correct allele of origin, SNVs annotations between mouse species were fetched from the Mouse Genome Project portal (57). SNVs were injected into the UCSC *Mus musculus* reference genome distributed through the Bioconductor (58, 59) package BSgenome.Mmusculus.UCSC.mm10 (60) using the function *qAlign* provided with the Bioconductor package QuasR (61). This resulted in two separate genomes to which reads were competitively aligned using the QuasR function *qAlign* with alignment parameters “-e 70 -X 1000 -k 2 –best –strata”. The best alignment determined the allelic assignment for each read. Finally, duplicated reads were identified and discarded using the Picard tool *MarkDuplicates* v2.15.0 (68).

### Targeted SMF at the Ectopic Locus Data Pre-processing

Amplicon SMF sequencing reads were trimmed for Illumina adapters and low-quality bases using TrimGalore v0.6.7. Reads shorter than 20bp after trimming were discarded. Afterward, a custom R script was used to trim the plasmid backbone from the reads. Pre-processed reads were mapped to the bisulfite indexed *Mus musculus* reference genome using the QuasR function *qAlign* with alignment parameters “-e 70 -X 1000 -k 2 –best –strata”. Duplicated reads were detected by the methylation pattern of cytosines and removed using custom functions.

### Definition of SMF Methylation Calling Windows

3,287,189 genomic tiles were designed to cover the genomic regions enriched during the SMF bait capture step. Each genomic tile spanned 80bp and overlapped the successive tile by 40bp. Iterating over large numbers of methylation calling windows, even when done in parallel, results in prohibitive computation times. Therefore, we designed large methylation calling windows encompassing multiple genomic tiles. To this end, the function *Create_MethylationCallingWindows* from the Single-MoleculeFootprinting package was used with parameters *fix*.*window*.*size* set to TRUE and *max*.*window*.*size* set to 50.

### CRE Annotation with ChromHMM

A whole genome 12-states chromatin annotation for mESCs was obtained from (69). For simplicity, we selected and pooled states as CTCF (1_Insulator), intergenic (2_Intergenic), enhancer (4_Enhancer, 8_StrongEnhancer, 11_WeakEnhancer), and promoter (7_ActivePromoter).

### Number of TF Motifs at CREs

The number of TF motifs at CREs was defined using the *Arrange_TFBSs_clusters* function from the SingleMoleculeFootprinting Bioconductor package. A list of bound and measured motifs was passed to the *TFBSs* argument. The arguments *max_cluster_width* and *max_cluster_size* were set to 300 and 10, respectively.

### TFBS Loss of Function Annotation

For each TFBS, the relative change in PWM match score across alleles was computed as (*A − R*)*/R*. TFBSs were considered loss of function at the alternative allele if their relative change was lower than −0.5. Gain of function motifs (relative change greater than 1.5) were discarded due to the lack of ChIP-seq/-nexus data for validation. All the remaining TF motifs were considered unperturbed across alleles.

In some instances, the Bl6xCast and Bl6xSpret F1s presented the very same genetic variation. To avoid such redundancies, we retained only the TFBS from the F1 cross with the highest SMF coverage.

### PRO-seq Statistical Testing

For each TF motif and genotype, the PRO-seq signal covering the 200bp surrounding the motif was quantified using the *qCount* function of the QuasR package. Statistical testing was performed using the DESeq2 Bioconductor package (64).

### SMF - Single Locus Plots

SMF bulk plots were generated using the function *PlotAvgSMF* from the SingleMoleculeFootprinting package. *FootprintCharter* heatmaps were generated using custom functions. Molecules were sorted for plotting using the partitioning results from *FootprintCharter*.

### Statistical Testing and Correlation Coefficients

Testing for statistically significant differences in the distributions of chromatin accessibility frequency values, or deltas in PRO-seq log2 fold changes, was carried out using the R implementation of the Wilcoxon signed-rank test.

Correlation coefficients were computed using the R function *cor* with the argument *method* set to *pearson*.

### Data Visualization and Schematic Illustrations

Scatter-plots, boxplots, violin plots, density plots, volcano plots, and genomic tracks displayed throughout the manuscript were generated using the R package *ggplot2*. Heatmaps were generated using the *ComplexHeatmap* package (70). Schematic illustrations displayed throughout the manuscript were generated using *Biorender*.

### Scripting, Data Analysis, and High-Performance Computing

All the scripting and data analysis for this project was performed using R-4.2.2 (71) and *RStudio Pro 2023*.*12*.*0+369*.*pro3* (72). The pre-processing pipeline for SMF data was scripted using *Nextflow-22*.*10*.*6* (73). All high-performance computing was performed using the *SLURM* workload manager (74), either through direct bash scripting or through the R package *rslurm* (75).

## Acknowledgments

We would like to thank Michela Palamin, Till Schwämmle, Tineke Lenstra, Anthony Mathelier, and Svetlana Dodonova for feedback on the manuscript. We thank Alexandra Maritz for help in generating F1 mESC lines. We are grateful to Ralph Grand for technical advice on targeted SMF at the ectopic locus. We are also grateful to the DKFZ mouse facility for crosses and deriving the F1 mESCs. Research in the laboratory of A.R.K. is supported by core funding from the EMBL, Deutsche Forschungsgemeinschaft (KR 5247/1-2, KR 5247/1-3) and the ERC (TFCoop-101125530).

We would like to thank Dirk Schübeler and Noa Gil for independently confirming the genomic location of the landing pad. The salaries of G.B. and V.B. were supported by the Deutsche Forschungsgemeinschaft (KR 5247/1-2). The generation of the Bl6xCast F1 mESC line was supported by the ERC (CTCFStableGenome-788937). The authors are grateful to GBCS and GeneCore at EMBL for support in sequencing data generation and management.

## Contributions

V.B. and G.B. co-led and contributed equally to the work. A.R.K. conceptualised this project with the support of G.B. and V.B. A.R.K. wrote the manuscript together with G.B. and V.B., and input from J.B.Z. G.B. conceptualised and implemented *FootprintCharter*. G.B. performed data analysis. V.B. established the reporter assay, generated the ectopic SMF data, and performed the analysis with support from G.B. R.K. generated the data for F1 cell lines. V.B. generated the data with support from L.M.P. for the SOX2-degron and

R.K. for the p300 inhibition experiment. K.C. generated the PRO-seq data. C.D., T.H. derived the Bl6xSpret F1 mESC lines. D.T.O., M.S., and R.K. generated the Bl6xCast F1 mESC line. A.R.K. acquired the funding. A.R.K and J.B.Z. supervised the work. All authors discussed the results and approved the manuscript.

## Conflict of Interest

The authors declare that they have no conflict of interest.

## Additional Information

Correspondence and requests for materials should be addressed to J.Z. (*Judith*.*zaugg@embl*.*de*) and A.R.K. (*arnaud*.*krebs@embl*.*de*).

## Code Availability

*FootprintCharter* is distributed with the devel version of the Bioconductor package *SingleMoleculeFoot-printing* at https://github.com/Krebslabrep/SingleMoleculeFootprinting/tree/dev. The code used to produce the figures in this manuscript is available at https://github.com/Krebslabrep/TF-chromatin.git.

## Supplementary Figures

**Supplementary Figure 1.**
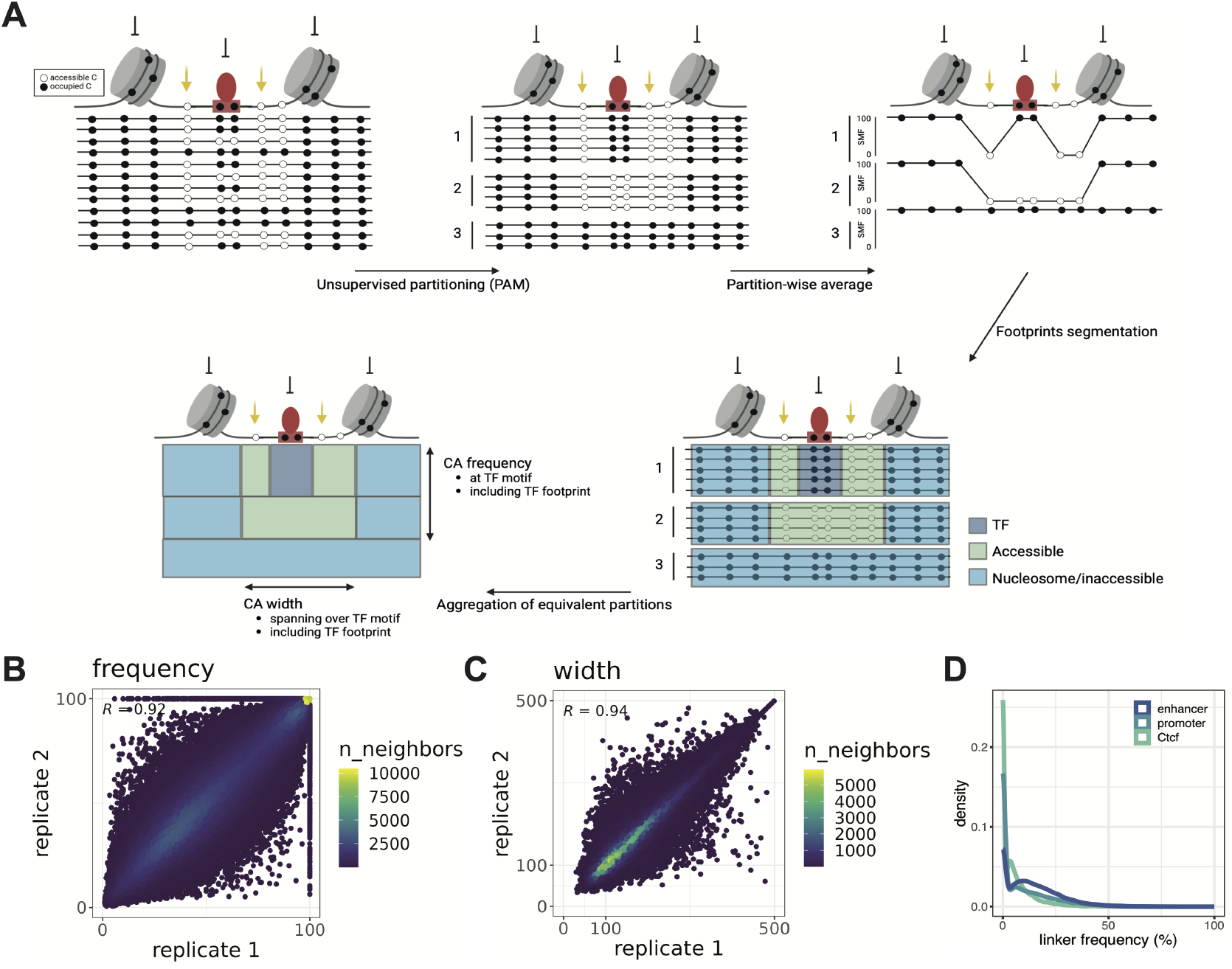
**(A)** *FootprintCharter* performs unsupervised molecular partitioning using the k-medoids algorithm. The SMF bulk signal is computed separately for each partition and used to detect TFs and nucleosome footprints by frequency and width. Such footprint detection is performed agnostic of TF motif annotations. These TF annotations are then used to aggregate biologically equivalent partitions to compute the frequency and average width of chromatin accessibility at TF motifs. **(B)** Scatterplot showing the replicate reproducibility of the chromatin accessibility frequency metric as calculated by FootprintCharter. The color scale indicates the number of neighboring dots, which shows the degree of overplotting. Pearson’s correlation coefficient is annotated. **(C)** Scatterplot showing the replicate reproducibility of the chromatin accessibility width metric as calculated by *FootprintCharter*. The color scale indicates the number of neighboring dots, showing the degree of overplotting. Pearson’s correlation coefficient is annotated. **(D)** Density plot showing the distribution of frequencies of linker-length DNA accessibility across active regulatory elements.

**Supplementary Figure 2.**
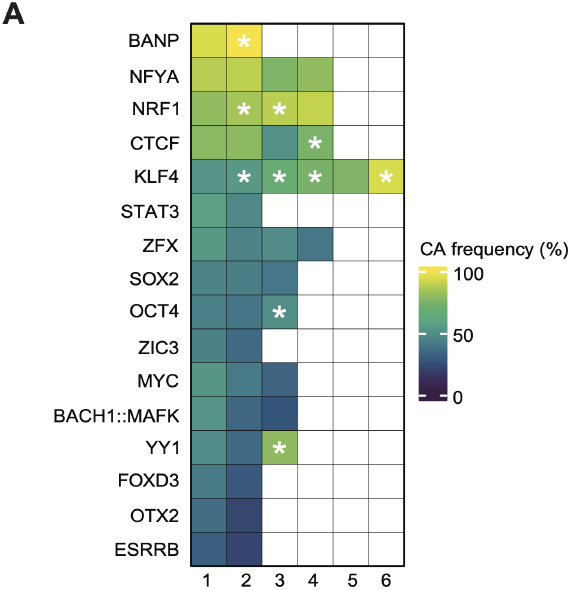
**(A)** Heatmap showing the median values of chromatin accessibility frequencies at active molecules associated with an increasing number of motifs (columns) for various TFs (rows). Cells annotated with stars indicate higher frequencies as compared to CREs with one motif less for the same TF (Wilcoxon test, *p <* 0.05).

**Supplementary Figure 3.**
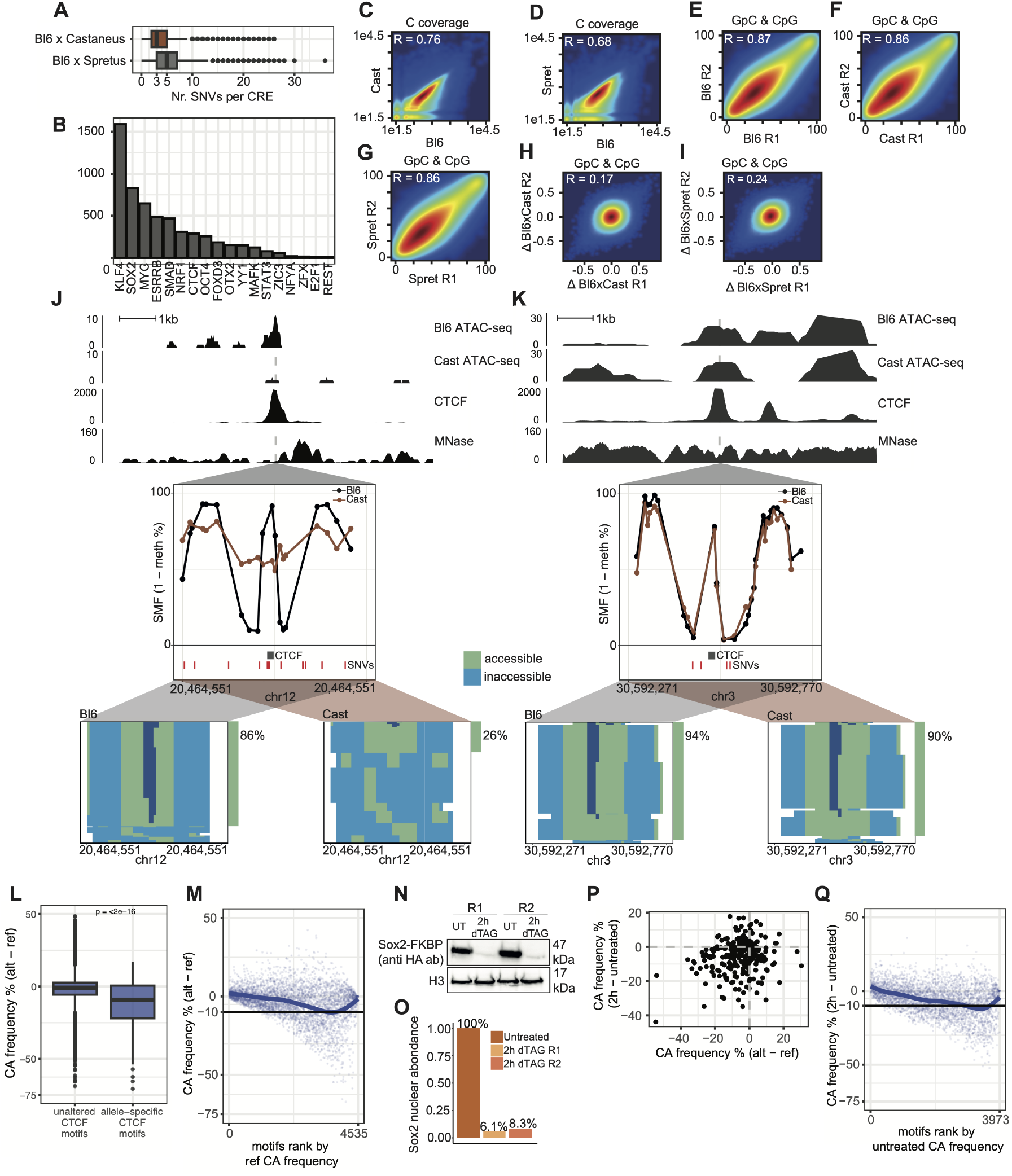
**(A)** Boxplots showing the distributions of number of SNVs per CRE in the Bl6xCast (brown) and Bl6xSpret (grey) F1s. **(B)** Bar-plot showing the number of TF motifs with allele-specific affinities across F1s. **(C-D)** Distribution of cytosine coverage across alleles for the Bl6xCast (C) and Bl6xSpret (D) F1s. The Pearson correlation coefficient *R* is annotated. **(E-G)** Distributions of methylation rates (%) for GpCs and CpGs across biological replicates for the Bl6 (E), Castaneus (F), and Spretus (G) alleles. The Pearson correlation coefficient *R* is annotated. **(H-I)** Distributions of methylation rate deltas across biological replicates for the Bl6xCast (H) and Bl6xSpret (I) F1s. The Pearson correlation coefficient *R* is annotated. **(J)** Example CTCF motif with allele-specific affinity between the Bl6 and Castaneus alleles associated with a 60% reduction in chromatin accessibility frequency. **(K)** Example CTCF motif with no difference in affinity between the Bl6 and Castaneus alleles and no change in chromatin accessibility frequency. **(L)** Boxplot of chromatin accessibility (CA) frequencies between alleles (alt - ref) at unchanged and allele-specific CTCF motifs across the mouse genome. The Wilcoxon test p-value is annotated. **(M)** Scatterplot of chromatin accessibility (CA) frequencies between alleles (alt - ref), ranked by accessibility frequencies on the reference allele. **(N-O)** Western blot reporting the loss in SOX2 protein abundance upon 2h dTAG treatment (N), quantified in (O). **(P)** Scatterplot comparing the chromatin accessibility (CA) frequencies in the dTAG experiment (2h dTAG - untreated) and in the F1 mESCs (alt - ref). **(Q)** Scatterplot of chromatin accessibility (CA) frequencies between alleles (2h dTAG - untreated) at accessible molecules, ranked by accessibility frequencies in the untreated condition.

**Supplementary Figure 4.**
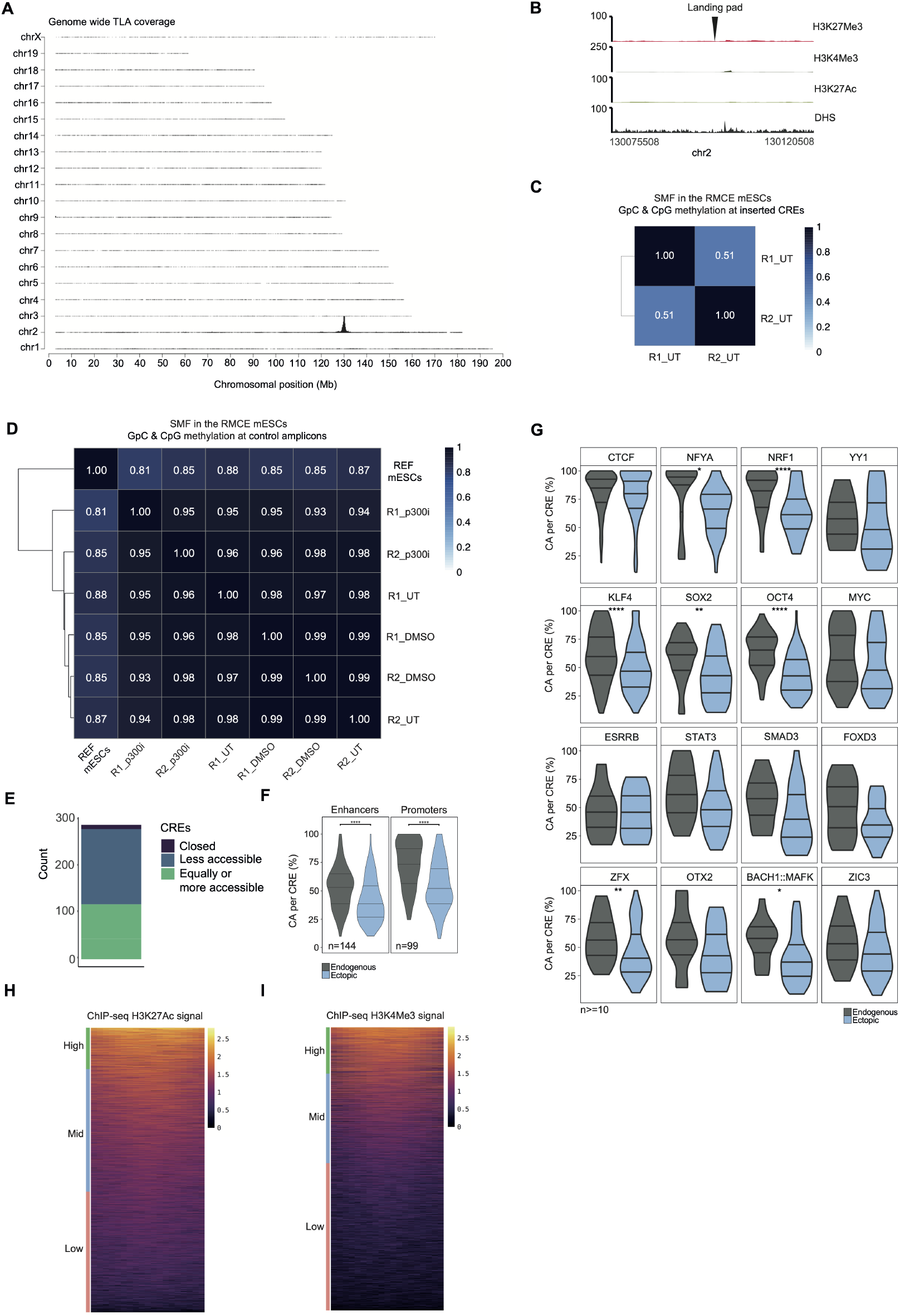
**(A)** Targeted Locus Amplification (TLA) signal across the mouse genome. The chromosomes and the chromosomal position are shown. The landing pad is inserted on chr2:130,098,005-130,098,011. **(B)** Genome browser tracks displaying levels of H3K4Me3 and H3K27Ac and chromatin accessibility at the landing pad at the neutral ectopic locus (chr2: 130,075,508- 130,120,508 – 45kb window of observation). **(C)** Pearson correlation heatmap of the methylation rates of cytosines in different contexts (i.e., CpG and GpC) across biological replicates. Methylation rates were measured by SMF at the genomic regions inserted at the ectopic site in untreated (UT), steady state condition. The dendrogram (left) shows hierarchical clustering of methylation signal, using complete-linkage method. The sample order reflects their similarity in methylation signal, with branch lengths indicating the relative distance between clusters. The Pearson correlation coefficient *R* is annotated. **(D)**Pearson correlation heatmap of the methylation rates of cytosines in different contexts (i.e., CpG and GpC) across biological replicates in untreated (UT), DMSO or p300 inhibition (p300i) conditions. Methylation rates were measured by SMF at the 4 endogenous control loci in the DNA methyltransferase triple-knockout (DNMT TKO, reference mESCs) and in the RMCE TC-1 DNMT TKO cells. Same representation as (C). **(E)**Stacked bar plot showing the proportion of DNA fragments that are less or equally accessible at the ectopic locus compared to their endogenous counterpart. **(F)**Violin plots showing the frequency of chromatin accessibility at enhancers (left) and promoters (right) when measured at the endogenous loci (black) and at the ectopic locus (blue). **(G)**Violin plots showing the frequency of chromatin accessibility at CREs when measured at the endogenous loci (black) and at the ectopic locus (blue) as a function of the presence of specific TF motifs within the DNA fragments, as defined by ChIP-seq and motif annotation (see Methods: TFBS annotation). **(H)**H3K27Ac ChIP signal, normalized per ChIP input and spike-in ranked from high to low. The H3K27Ac signal is calculated in 2kb windows covering Transcription factor binding sites (TFBSs) in the genomic regions enriched during the SMF bait capture step (see Methods for TFBS annotation). Shown is the log10 of the H3K27Ac coverage. Classes were defined based on the enrichment pattern, as: low (<100), mid (between 100 and 500), and high (>500). **(I)**H3K4Me3 ChIP signal, normalized per ChIP input and spike-in ranked from high to low. The H3K4Me3 signal is calculated in 2kb windows covering Transcription factor binding sites (TFBSs) in the genomic regions enriched during the SMF bait capture step (see Methods for TFBS annotation). Shown is the log10 of the H3K4Me3 coverage. Classes were defined based on the enrichment pattern, as: low (<50), mid (between 50 and 300), and high (>300).

**Supplementary Figure 5.**
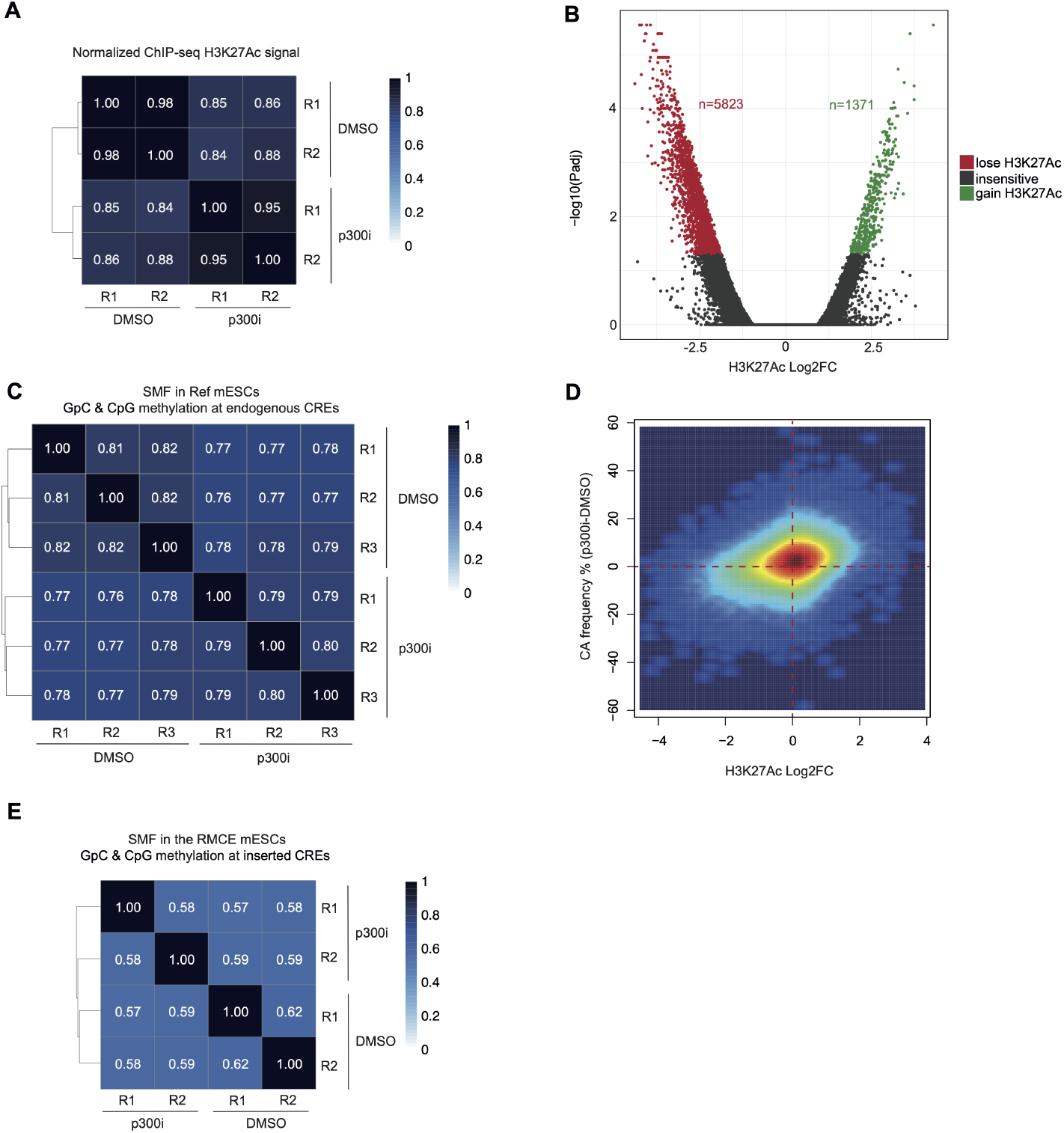
**(A)** Pearson correlation heatmap of H3K27Ac ChIP-seq signal across biological replicates in DMSO and p300i treatment, normalized per input and spike-in (see Methods for normalization steps). H3K27Ac signal is calculated in 2kb windows covering the genomic regions enriched during the SMF bait capture step. The dendrogram (left) shows hierarchical clustering of H3K27Ac signal, using complete-linkage method. The sample order reflects their similarity in H3K27Ac signal binding, with branch lengths indicating the relative distance between clusters. The Pearson correlation coefficient *R* is annotated. **(B)** Volcano plot showing the H3K27Ac log_2_ fold change (Log_2_FC, x-axis) and −log_10_ p-value (y-axis) probing the changes of H3K27Ac enrichment upon p300 inhibition at endogenous loci. The log_2_ fold change of H3K27Ac was calculated using the DESeq2 package, providing H3K27Ac counts in DMSO or p300i treatment replicates, after normalizing per input and spike-in (see Methods for normalization steps). **(C)** Pearson correlation heatmap of the methylation rates of cytosines in different contexts (i.e., CpG and GpC) across biological replicates in DMSO or p300i treatment. Methylation rates were measured by SMF at the genomic regions enriched during the SMF bait capture step, in DNMT TKO mESCs (Ref). Same representation as (A). **(D)** Scatterplot showing the distribution of chromatin accessibility (CA, p300i-DMSO, y-axis) frequencies measured at endogenous loci as a function of the H3K27Ac log_2_ fold change (Log_2_FC, x-axis) upon p300 inhibition at the same loci. **(E)** Pearson correlation heatmap of the methylation rates of cytosines in different contexts (i.e., CpG and GpC) across biological replicates in DMSO or p300i treatment. Methylation rates were measured by SMF at the genomic regions inserted at the ectopic site. Same representation as (A).

### Supplementary Tables

**Supplementary Table 2.**
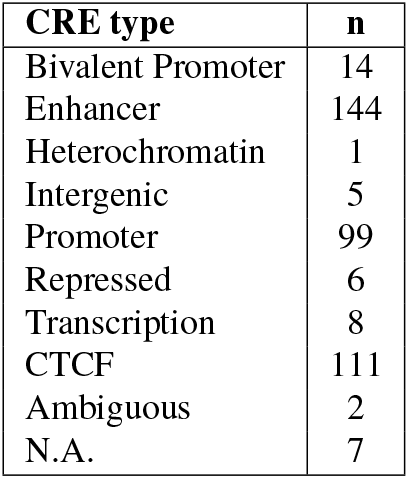
Composition of fragments with sufficient coverage retrieved from the library inserted at the ectopic locus.

**Supplementary Table 3.** Details of the performed assays and corresponding cell lines.

| Assay | Cell line |
| --- | --- |
| SMF-bait capture | F1 mESCs |
| PRO-seq | F1 mESCs |
| targeted-SMF | TC-1 DNMT TKO mESCs (RMCE) (untreated, DMSO, p300 inhibition) |
| SMF-bait capture | SOX2 degron mESCs |
| SMF-bait capture | 159 DNMT TKO mESCs (DMSO, p300 inhibition) |
| ChIP-seq | 159 DNMT TKO mESCs (DMSO, p300 inhibition) |

## Notes

### Competing Interest Statement

The authors have declared no competing interest.

